# WHEN MICROSCALE TRANSPORT CONTROLS MACROSCALE TISSUE FREEZING: A PARAMETRIC REGIME ANALYSIS

**DOI:** 10.64898/2026.09.26.754647

**Authors:** R.V. Devireddy

**Affiliations:** Department of Mechanical Engineering, Louisiana State University, Baton Rouge, LA 70809 USA

**Keywords:** cryopreservation, enthalpy/apparent-heat-capacity finite-volume model, osmotic dehydration, tissue damage kinetics, parametric sensitivity analysis, intracellular ice formation

## Abstract

Devireddy et al. [1] previously showed that an enthalpy-based, macroscale model simulated tissue freezing histories are in close agreement with a coupled model incorporating cellular water transport and intracellular ice formation (IIF). That comparison, however, was performed for a limited set of tissue geometries, cooling conditions, and biophysical parameters. Here, we revisit that conclusion by reimplementing the published model from its governing equations and systematically expanding the parameter space to determine when microscale processes alter the predicted macroscale freezing response. We incorporate both surface-catalyzed nucleation (SCN) and volume-catalyzed nucleation (VCN) of intracellular ice with cellular osmotic dehydration and examine a broad range of tissue dimensions, cooling rates, convective boundary conditions, membrane permeabilities, cell sizes, activation energies, intracellular water fractions, and nucleation rates. Linearization of the cellular water-transport equation about osmotic equilibrium yields an osmotic relaxation time (*τ*_*osm*_) that can be compared with the local residence time of tissue undergoing phase change (*τ*_*res*_). Their ratio defines a dimensionless group, ∏, that organizes the transition between regimes in which the uncoupled enthalpy formulation and the coupled micro–macroscale formulation produce similar or substantially different results. Across the one-dimensional parameter space examined here, the two models remain within a prescribed thermal-error bound for ∏ below approximately 0.1, whereas the discrepancy increases systematically above this threshold. Model divergence also depends on intracellular ice nucleation, i.e., when nucleation is sufficiently slow, cellular water transport can become rate-limiting for latent-heat release. Conversely, sufficiently rapid intracellular ice formation reduces the macroscopic discrepancy by providing an additional pathway for latent-heat release. Multidimensional simulations further show that geometry alone can move a nominally conventional freezing protocol into the coupled regime. In particular, multidirectional cooling can reduce the local residence time sufficiently to produce substantial differences in predicted frozen volume even when the corresponding thermal histories appear similar. Finally, we formulate an inverse approach for estimating IIF parameters from tissue-scale thermal histories. Synthetic simulations indicate that parameter recovery is feasible within a finite range of cooling conditions, providing a testable strategy for future experimental measurements. Together, these results provide a mechanistic criterion for deciding when microscale biophysical processes must be retained in macroscale tissue-freezing models.

Graphical Abstract
The design space for tissue freezing is illustrated as a function of a non-dimensional grouping parameter, distinguishing between the uncoupled (equilibrium) model and the coupled micro–macro-scale model. Cooling conditions corresponding to grouping values below 0.1 can be adequately described using the uncoupled model, whereas values above 0.1 require the coupled micro–macro-scale tissue-freezing model. The critical value of 0.1 is established based on a maximum acceptable temperature difference of 3 K between the two models. The operating point reported in the 2002 study lies approximately two orders of magnitude within the uncoupled regime and is indicated by a star. In contrast, an 8 mm cube cooled on all six faces falls within the coupled regime and is represented by a red dot.

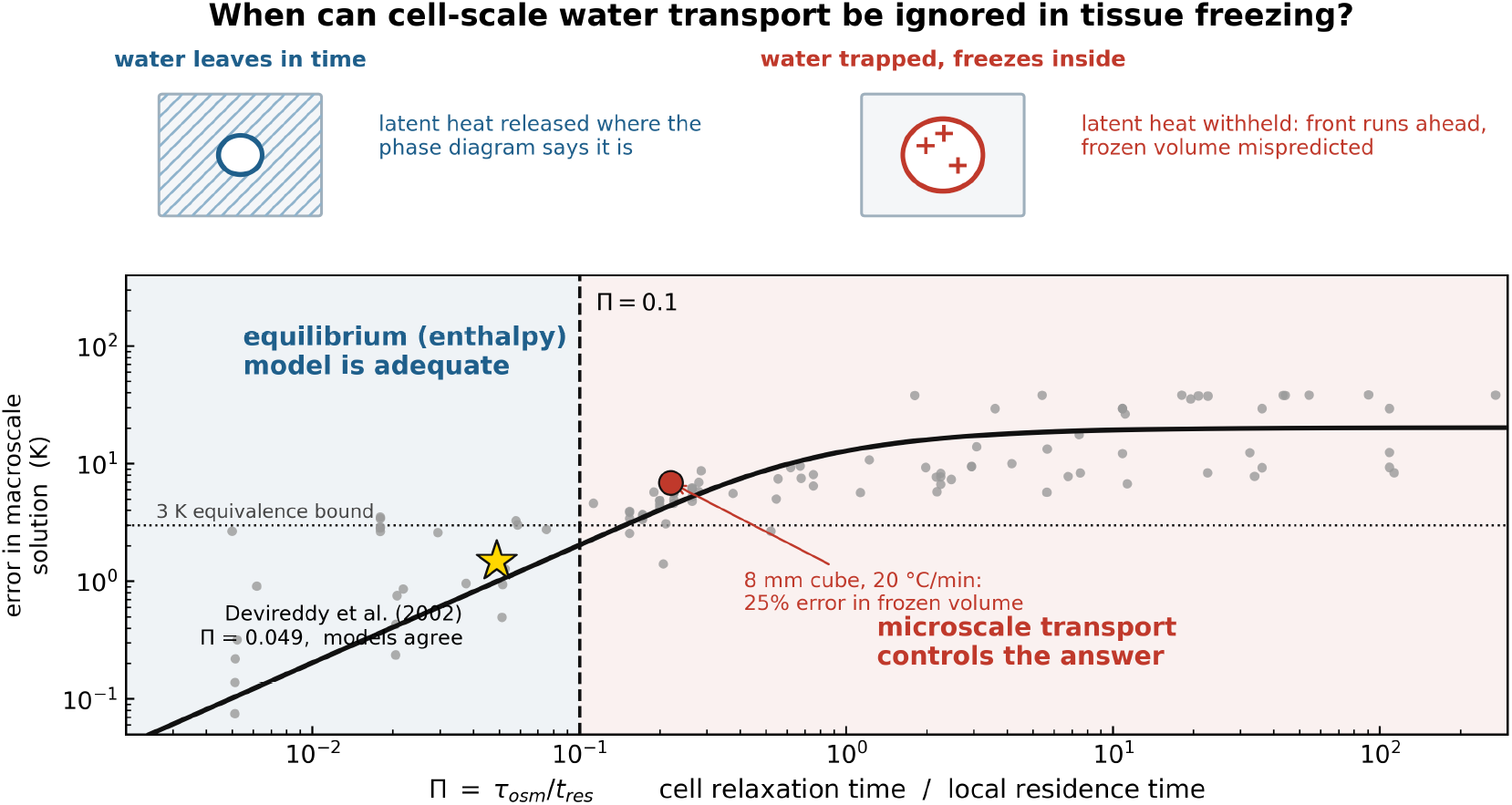

**Nomenclature^1^:** 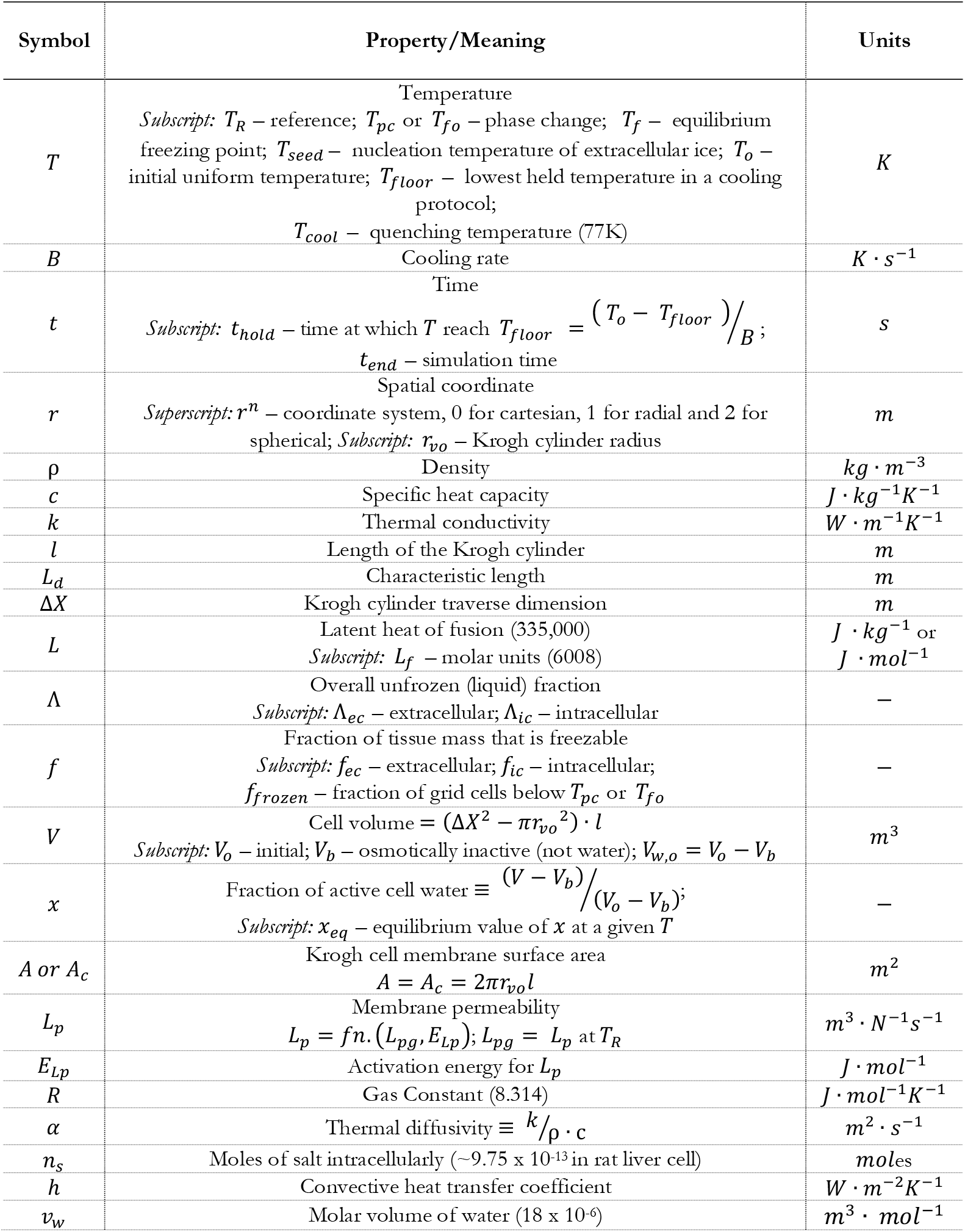

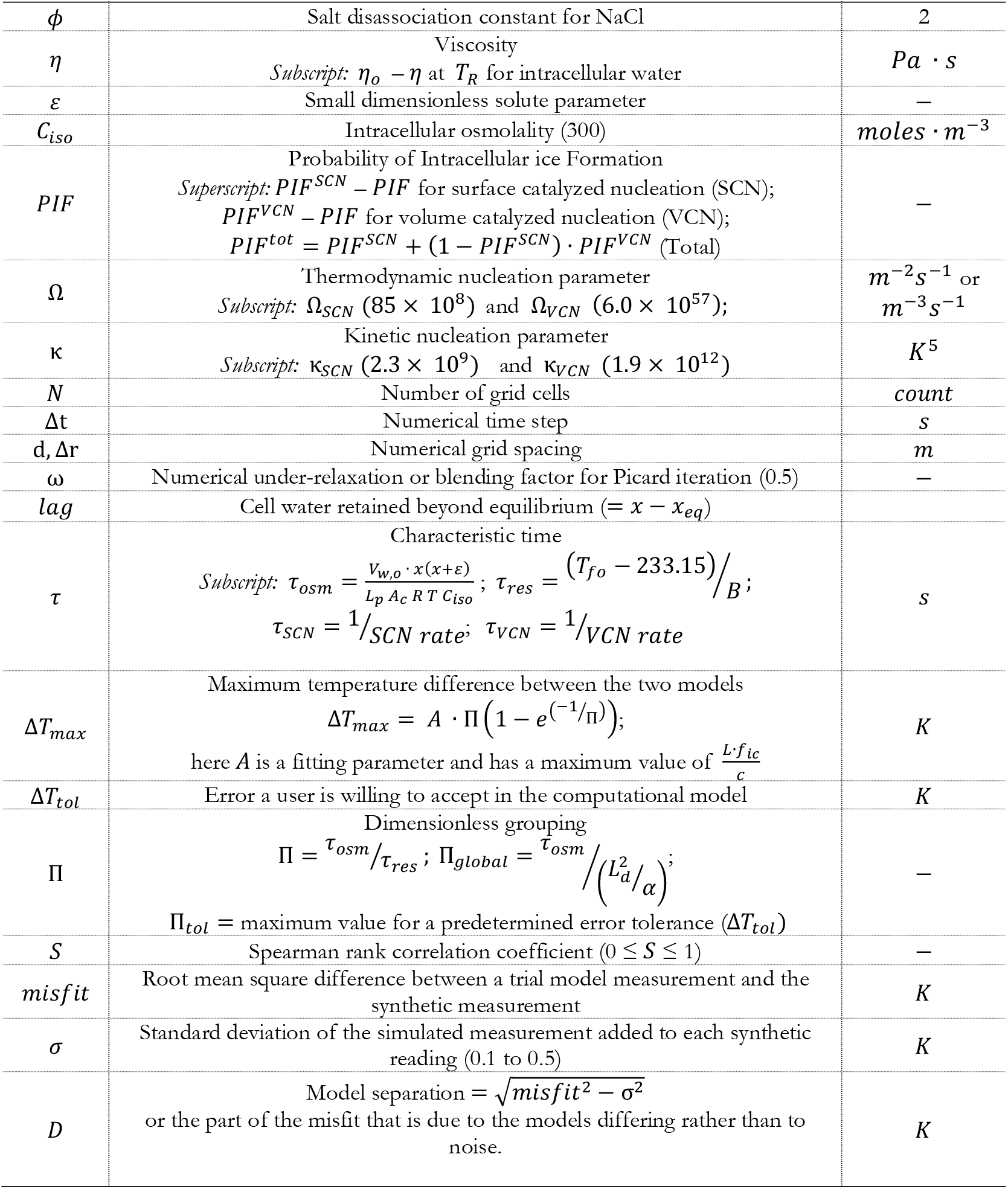

## INTRODUCTION

The process of freezing biological tissue is an active area of interest for cryobiologists intent on optimizing cryopreservation (the storage of biological systems at low temperatures) or cryosurgery (the destruction of tumorous tissue using low temperatures). The interested reader is referred to several historical and recent articles on the topic [2–16]. Two principal biophysical processes govern cellular responses during tissue freezing: cellular dehydration and intracellular ice formation (IIF). During dehydration, intracellular water leaves the cell and contributes to extracellular ice formation or alternatively, intracellular water can nucleate and form intracellular ice [4]. Both processes can compromise cell viability and function. Mazur’s two-factor hypothesis describes two limiting regimes, i.e., at low cooling rates, injury is associated primarily with solute concentration effects caused by cellular dehydration, whereas at high cooling rates, injury is associated primarily with intracellular ice formation (IIF). The exact magnitude of the ‘‘low’’ and ‘‘high’’ cooling rates are specific to a cell type and is dependent on the biophysical water transport and IIF parameters [4,7–10,17]. Both water transport and IIF have been extensively measured and mathematically modeled in isolated cells [4,7,8,10,17–34]. Water transport has also been measured and modeled in tissues [34–43]. In contrast, quantitative methods for measuring IIF kinetics directly in intact tissues remain limited. In addition, numerous computational methods and models have been used to elucidate the process of freezing in tissues, as evidenced by the volume of published literature [1,8,10,23–25,34,36–68].

To accurately model the freezing processes in tissues it is important to assess the conditions under which the microscale biophysical phenomena (water transport and IIF) play a role in determining the macroscale tissue thermal history. The initial attempt at this was by Rubinsky and Pegg [69] when they proposed a mathematical model for tissue freezing in which a cell dehydration model and the phase diagram for isotonic saline, was used to govern the rate of phase change. Subsequently, Bischof and Rubinsky [70] incorporated a model for IIF, and solved the coupled thermal and biophysical model numerically for a one-dimensional cartesian coordinate system, using a front tracking method. Devireddy et al. [1] subsequently formulated the microscale processes as a temperature-and time-dependent latent-heat source term, Λ(*T, t*) and solved the resulting coupled heat-transfer problem in one spatial dimension. The analysis considered rat liver cooled at 5 °C/min and AT-1 tumor tissue cooled at 50 °C/min. It was restricted to two tissue types, specific geometries, and prescribed membrane-permeability and nucleation parameters, and included only surface-catalyzed nucleation (SCN). Volume-catalyzed nucleation (VCN), which becomes relevant at lower temperatures, was not included. Despite these restrictions, the coupled and uncoupled formulations produced similar freezing predictions for the cases examined. This result established an important baseline, but it did not determine the conditions under which the microscale processes become rate-limiting. Recent advancements in millimeter-scale, multi-directionally cooled and rapidly quenched protocols used in vitrification-based organ preservation and in engineered-tissue processing has made this question relevant and has not, to our knowledge, been systematically examined, and no criterion has been proposed for identifying, in advance, where the simplification stops holding [11–16].

Here, we revisit this question computationally. We reimplemented the coupled and uncoupled models developed in the earlier study [1] directly from their governing equations and verified the reimplementation against the original published results. We then inverted the design of the original study. Rather than evaluating whether cellular kinetics are important at a single operating point, we systematically explored a broad range of sample geometries, cooling protocols, and cell and tissue properties across one, two, and three spatial dimensions. This allowed us to identify the conditions under which microscale biophysical processes influence the macroscale tissue freezing problem. We present our analysis by deriving a single dimensionless group defined as the ratio of a cell’s osmotic relaxation time to the local residence time within the phase change window. This dimensionless group collapses the observed model error across all cases examined and provides a closed form explanation for why the original result holds under the conditions that were tested and where it begins to break down. Our analysis further identifies intracellular ice formation as a second thermally compensating pathway and demonstrates that routine multidimensional cooling protocols can cross the boundary between uncoupled and coupled behavior solely as a consequence of geometry. The resulting framework provides a quantitative criterion that can be calibrated to a user selected error tolerance and used to determine when the coupled microscale model developed by Devireddy et al. [1] is required and justified. In addition, our analysis enabled us to propose a methodology for measuring the intracellular ice formation parameter in tissues. This methodology will be fully developed and evaluated through complementary computational and experimental studies presented in a companion paper.

### GOVERNING EQUATIONS

Freezing in tissue is described by the coupled macroscale/microscale formulation which we adopt without modification [1]. Briefly, energy transport follows conduction with a latent-heat source as,

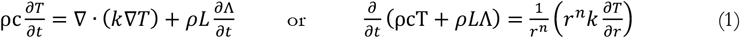

where, ρ is the density, c is the specific heat capacity, *k* is the thermal conductivity, Λ is the unfrozen water fraction, *T* is temperature, *t* is time, and *L* the latent heat of fusion of water. The value of n determines the coordinate system with n = 0 for cartesian, n = 1 for cylindrical and n = 2 for spherical coordinate systems. In tissues, freezable water is partitioned between extracellular and intracellular compartments as: Λ = *f*_*ec*_ · Λ_*ec*_ + *f*_*ic*_ · Λ_*ic*_, where *f*_*ec*_ and *f*_*ic*_ are the fractional masses of extracellular and intracellular water in the medium, respectively. The fractional mass is the mass of water divided by the total mass of medium. Assuming that biological tissue has a density equal to water, the fractional masses *f*_*ec*_ and *f*_*ic*_ can be approximated by the volume fractions of the extra and intracellular water respectively; the quantities *f*_*ec*_ and *f*_*ic*_ will therefore be referred to as fractional volumes in the remainder of the paper, since this is how they are calculated. In general, because of the presence of non-water components in biological tissue, the sum *f*_*ec*_ and *f*_*ic*_will be < 1. The extracellular latent heat fraction Λ_*ec*_ is defined using the binary phase diagram for a 150 mM NaCl aqueous solution, as given by [71–75]:

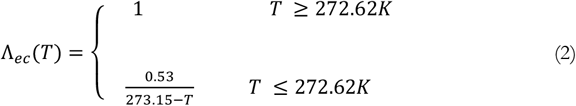

The intracellular latent heat fraction Λ_*ic*_ is an integral function of cell volume (*V*) and probability of intracellular ice formation (*PIF*) as [1]:

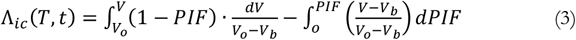

or by tracking both contributions over a small time step gives the incremental (step-by-step) update as:

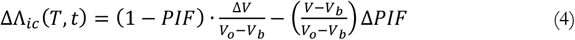

where, *V*_*o*_ is the initial cell volume (before freezing begins), and *V*_*b*_ is the volume of osmotically-inactive, i.e., non-water cell components. The first integral represents the contribution of intra-cellular water which freezes after being transported through the cell membrane to the extracellular space, and the second integral represents the contribution of water which freezes intracellularly. In order to evaluate these two integrals, both Δ*V* and Δ*PIF* must be determined during the freezing process. A full description of the water transport model to determine *V*(*T, t*) and the intracellular ice nucleation model to determine *PIF*(*T, t*) are described elsewhere [4,7,8,10,11,20,23–25,27,29,34–36,45–48,71–78] and will only be briefly described here.

The model to describe cellular dehydration or *water transport during freezing* in biological systems at slow cooling rates predicts the change of cell volume with decreasing temperature (after ice has formed outside an unfrozen cell) is:

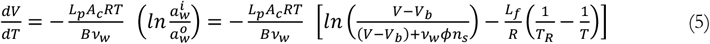

where *V* is the cell volume, *T* is the absolute temperature, *L*_*p*_ is the permeability of the membrane to water, *R* is the gas constant, *B* is the constant cooling rate, *A*_*c*_ is the effective membrane surface area available for water transport, *v*_*w*_ is the partial molar volume of water, and 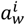 is the chemical potential of the intracellular fluid and 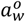 is the chemical potential of extracellular fluid. Quantitative use of the above thermodynamic model requires a more complete specification of the chemical activities of the extracellular and the intracellular solutions, along with the membrane water permeability [4,7,8,10,11,20,23–25,27,29]. The cell membrane permeability, *L*_*p*_, depends on two biophysical parameters following an Arrhenius temperature dependence as: 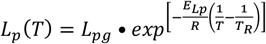, with *L*_*pg*_ denoted as the reference water permeability at the phase-change temperature of water, and *E*_*Lp*_ as the activation energy for the water transport process. Both the membrane permeability parameters (*L*_*pg*_, *E*_*Lp*_) are also functions of the type of CPA as well as the amount of CPA in the freezing medium and are also dependent on the presence or the absence of extracellular ice [4,7,8,10,11,20,23–25,27,29].

The model to describe the IIF states that for a thermodynamic system composed of identical biological cells, the total probability of intracellular ice formation (PIF^tot^) is a function of heterogeneous or surface-catalyzed nucleation (PIF^SCN^) and homogeneous or volume-catalyzed nucleation (PIF^VCN^) as [23–25,76–78]:

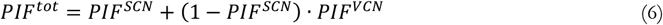

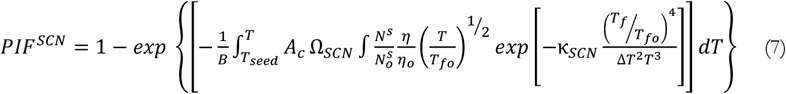

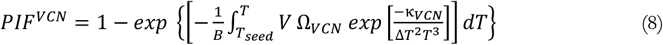

where the subscript ‘o’ refers to isotonic conditions; Δ*T* is the amount of supercooling (*T*− *T*_*f*_); all the other variables are as described elsewhere [*] and in the nomenclature section. Assuming that the nucleating site is the plasma membrane and that the number of solute molecules are much lesser than the total number of water molecules, the ratio of 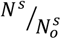 in Eqn. (7) can be written as *A*/*A*_*o*_ [23–25,76–78]. In order to evaluate Eqn. (7) during freezing of cells, both the concentration and temperature dependence of the viscosity, *η* is needed. However, in most studies, the dependence of viscosity on the concentration of cytoplasm is neglected and a power law dependence of water viscosity on temperature, T (K) is applied as: *η*(*T*) = 0.139 · {(*T*/225) − 1}^−1.64^ · 10^−3^ (Kg·m^−1^·s^−1^) and was also applied in the present study [76–87]. The volume catalyzed nucleation is assumed to occur spontaneously throughout the volume of the cytosol and there are currently no experimental techniques to quantify the volume catalyzed nucleation parameters (Ω_*VCN*_ and *κ*_*VCN*_) in biological cells. The parameters reported for physiologic cell-sized saline droplets, i.e. Ω_*VCN*_ = 6.0 × 10^57^ (m^−3^·s^−1^) and *κ*_*VCN*_ = 1.9 × 10^12^ (K^5^) are assumed to be representative of biological cells [87] and were used in the present study, as well. The fundamental difference between SCN and VCN are the way ice is nucleated either through a helper medium (foreign particles, or, importantly here, the cell membrane itself, that lower the energy barrier for nucleus/cluster formation) or heterogenous nucleation (SCN) and through homogeneous nucleation, which is the nucleation of ice in pure liquid with no helper surface at all (VCN). This difference in the nucleation process also implies that the temperature at which they occur are distinct and separate [23–25,76–78].

In summary, the thermodynamic state of the intracellular solution during freezing in the presence of extracellular ice can be characterized using a set of water-transport (*L*_*p*_) and heterogeneous (Ω_*SCN*_ and *κ*_*SCN*_) and homogeneous (Ω_*VCN*_ and *κ*_*VCN*_) ice nucleation parameters and these parameters enter the macroscale heat equation through the latent heat term, Λ(*T, t*), in Eqn. (1). Temperature-dependent thermophysical properties are those of water, and cellular geometry is represented by the Krogh cylinder (see full nomenclature listing for the exact values). Assumptions are those of the original formulation, i.e., homogeneous continuum for heat transfer, constant membrane area and osmotically inactive volume, ideal dilute solution, instantaneous extracellular equilibration, no cryoprotectant, no perfusion, and no mechanical coupling. The baseline membrane permeability (water transport) parameters (*L*_*pg*_, *E*_*Lp*_) as well as the osmotically inactive cell volume (*V*_*b*_) for rat liver and AT-1 tumor tissue were taken from published literature [34,36,88]. The baseline SCN values are those of isolated cells [77] rather than tissues, as these values are not available in literature. Note that replacing Λ_*ic*_(*T, t*) with Λ_*ec*_(*T*) eliminates the dependence of the tissue freezing problem on the micro-scale (water transport and IIF) phenomena and is denoted as the uncoupled model (or the classical enthalpy based formulation of tissue freezing).

### NUMERICAL METHODS

#### One dimension

We utilize a cell-centered finite volume in Cartesian and cylindrical coordinates, with harmonic-mean interface conductivities to conserve flux across the sharp conductivity jump at the front [89]. Time integration is fully implicit and the tridiagonal system is solved directly. Latent heat is treated by an apparent heat capacity, 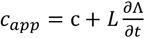, in which the derivative is evaluated as a secant slope over the time step [1]. The secant form is required rather than preferred, since for the coupled model Λ is history-dependent and has no analytic temperature derivative. The secant form is the only construction that treats both variants identically and conserves latent heat exactly over a step. The resulting nonlinearity is resolved by Picard iteration [90–92] under relaxation (with a blending factor, ω = 0.5), with convergence tested on the Λ field, max |Λ^(*m*)^ − Λ^(*m*−1)^| < *tol* < 10^−5^.

#### Two and three dimensions

The well-established Douglas–Rachford ADI with boundary sources entering once in the first directional sweep was utilized. All tridiagonal systems along a direction are solved simultaneously by a vectorized Thomas algorithm, which makes 2D/3D tractable while a sparse factorization per Picard iteration is not [93–100]. Phase change, relaxation and the convergence test are identical to the 1D scheme, and all constitutive functions are imported from the 1D module so that the three dimensionalities share one implementation. An explicit enthalpy scheme was evaluated first and rejected and at the grid spacings required here the stability limit gives: Δt ≈ 10^−5^secs.

#### Front tracking

The freezing front is defined as the T = *T*_*ph*_ isotherm while the trailing edge is marked by the Λ = 0.005 contour, where the freezable water is essentially exhausted. The two together bound the mushy zone and both are located by linear interpolation between adjacent nodes. The classical node-index detection is unusable in this scenario because of the mushy-zone (partially frozen) making the temperature field plateau near the freezing-phase change front and consequently, even a sub-millikelvin change displaces the front by several nodes [100–105]. All interface positions and front-arrival errors reported in this study refer to the leading edge while the trailing edge is recorded only for completeness and not used or reported.

#### Parametric design

The parameter space of Table 1 was explored in 124 one-dimensional cases organized in six families as sample thickness × cooling rate under a programmed ramp (36 cases); sample thickness × convective film coefficient under a quench (16); membrane permeability × cell size at a moderate and at a severe protocol (20 cases each or 40); activation energy × intracellular water fraction (20); and nucleation rate at three protocols (12). A further 42 cases mapped nucleation rate against protocol severity on a regular grid. They are the nucleation-window sweep: 6 protocols × 7 nucleation-rate multipliers (0, 10^−3^, 10^−2^, 10^−1^, 1, 10, 100 for Ω_*o*_). The protocols are the 2 mm slab at 2, 20, 60 and 200 °C/min, the 1 mm quench at h = 250*W* · *m*^−2^*K*^−1^, and the 0.5 mm quench at h = 1000 *W* · *m*^−2^*K*^−1^. The VCN constants were not varied as they are universal rather than tissue-specific. They do not appear in Table 1 because that table bins the parametric sweep by ∏ and these 42 runs deliberately hold ∏ fixed while varying only the nucleation rate. Each shares its Π with the parent protocol already counted in Table 1.The intra-cellular water fraction was tested at 0.25, 0.44, 0.589 and 0.62 (a planned 0.80 was discarded as that would have required a negative extracellular fraction, a physical impossibility). Cell size was scaled linearly, with *V*_*o*_ ∝ *s*^3^ and *A*_*c*_ ∝ *s*^2^ and the intracellular water fraction was varied with *f*_*ec*_ adjusted to hold *f*_*ec*_ + *f*_*ic*_ constant, so that total tissue latent heat is preserved. The programmed-ramp cases impose a surface temperature falling at a fixed rate to a hold, with the opposite face adiabatic (the symmetry plane of a slab cooled on both sides). Quench cases impose a convective condition into liquid nitrogen with h = 100 to 1000 W · m^−2^ · K^−1^, spanning film boiling to forced flow. Six multidimensional cases were also run: an axisymmetric cryosurgical probe (which, because the probe spans the full domain height with insulated ends, yields a solution uniform along the axis and is effectively radial), a tissue block cooled on two orthogonal faces, a vitrification carrier quenched on two faces, and three cubes cooled on all six faces (octant symmetry, exact for this configuration). Geometries, grids and boundary conditions are listed in Table 2.

**Table 1:** The parametric range explored in the present study vs. the earlier study of Devireddy et al. [1].

| Model Input | Range explored (Devireddy et al. 2002) | Range explored (current study) |
| --- | --- | --- |
| $d$ | 80 mm only | 0.25 to 80 mm |
| $B$ | 5 °C/min (rat liver) and<br>50 °C/min (AT-1 tumor) | 0.5 to 200 °C/min |
| $h$ | Not explored (Dirichlet BC only) | 100 to 1000 $W \cdot m^{-2}K^{-1}$ |
| $L_{pg}$ | 1x rat liver ($3 \times 10^{-13} m^3 N^{-1} s^{-1}$) and<br>1x AT-1 tumor ($0.5 \times 10^{-13} m^3 \cdot N^{-1} s^{-1}$) | 0.02x to 3x rat liver |
| $E_{lp}$ | 265 kJ·mol<sup>-1</sup> (rat liver) and<br>96.3 kJ·mol<sup>-1</sup> (AT-1 tumor) | 60 to 350 kJ·mol<sup>-1</sup> |
| $f_{ic}$ | 0.589 (rat liver) and 0.44 (AT-1 tumor) | 0.25 to 0.62<br>(at fixed total of 0.6265) |
| $\Omega_{scn}/\Omega_o$ | 1x only | 0 to 100x |
| $VCN$ | Off (SCN only) or not included | Off (SCN only) and<br>on (SCN + VCN) all cases |
| Spatial dimension | 1D only | 1D, 2D axisymmetric,<br>2D cartesian and 3D |
| Cooled faces | 1 | 1, 2, 3 (of a symmetric cell) |
| Krogh cylinder<br>dimensions | 1x rat liver<br>($\Delta X = 22 \mu m$; $r_{v,o} = 3.8 \mu m$; $l = 11.4 \mu m$) | 0.5x to 10x (rat liver) |
| $\tau_{osm}$ | 1x rat liver (24.3 s) and<br>1x AT-1 tumor (30.7 s) | 0.16x to 500x (rat liver) or<br>4.05 to 1.21 x 10<sup>4</sup> secs |
| $\tau_{res}$ | 1x rat liver (495 s) and<br>1x AT-1 tumor (700 s) | 0.00054x to 10x (rat liver) or<br>0.268 to 4854 s |
| $\Pi$ | 0.0491 (rat liver) and 0.0439 (AT-1 tumor) | 0.1x to 110,000x (rat liver) or<br>0.005 to 5422 |
| $\rho$ | 999 (unfrozen) and 921 $kg \cdot m^{-3}$ (frozen) | |
| $c$ | 4200 (unfrozen) and 7.16T+138 $J \cdot kg^{-1}K^{-1}$ (frozen) | |
| $\kappa$ | 0.6 (unfrozen) and 2.24 + 0.005975 (273 - T)<sup>1.156</sup> $W \cdot m^{-1}K^{-1}$ (frozen) | |

**Table 2:** Geometry, grid, boundary conditions and numerical settings for the six two- and three-dimensional cases. All use rat liver properties. Faces not listed as cooled are adiabatic. Octant and quarter domains are exact by symmetry.

| Geometry | Protocol | Grid | $\Pi$ | $\Delta T_{max}$<br>(SCN) | $\Delta T_{max}$<br>(SCN + VCN) | Frozen volume<br>gap (percentage<br>points) |
| --- | --- | --- | --- | --- | --- | --- |
| 2D Axisym.<br>(z-uniform;<br>radial) | 3mm cryoprobe; 50<br>°C/min to -160 °C | 110 x 110 | 0.33 | 4.7 | 4.7 | 1.8 |
| 2D Car. | 4mm block; 60 °C/min,<br>two faces | 100 x 100 | 0.67 | 7.4 | 7.4 | 38.3 |
| | 1 mm block;<br>$LN_2$ quench,<br>$h = 500 \text{ W} \cdot \text{m}^{-2} \text{K}^{-1}$ | 80 x 80 | 97.12 | 71.3 | 34.6 | 39.6 |
| 3D (6 faces<br>cooled) | 8 mm cube; 20 °C/min | 40 x 40 x 40 | 0.22 | 7.0 | 7.0 | 25.4 |
|  | 4 mm cube; 60 °C/min | 40 x 40 x 40 | 0.66 | 5.6 | 5.6 | 32.8 |
| | 1 mm cube;<br>$LN_2$ quench,<br>$h = 500 \text{ W} \cdot \text{m}^{-2} \text{K}^{-1}$ | 36 x 36 x 36 | 131.25 | 70.8 | 44.2 | 31.7 |

Every test case described was run twice, once with the uncoupled (enthalpy) model and once with the coupled model and on identical grids, time steps and solver settings, so all differences are attributable to the constitutive treatment of latent heat rather than discretization. However, a comparison of two formulations is only as good as the convergence of the stiffer one. We found that Δ*T*_*max*_ is insensitive to mesh refinement, changing by less than 0.1% between 200, 400 and 800 cells, and insensitive to the time step once the iteration is converged, changing by about 1% across an eight-fold range of Δ*t*. It is, however, sensitive to the Picard tolerance. At a tolerance of 10^−8^ with 200 iterations the residual iteration error does not cancel between the two models, because the coupled source term is stiffer than the equilibrium one, and individual cases moved by up to a factor of five in either direction when the iteration was tightened. All results reported here therefore use a tolerance of 10^−8^ with up to 200 iterations, verified case by case against one halving of the time step. Fifteen of the 124 one-dimensional cases still retain a residual time-step sensitivity of 3 to 7% and are reported with that uncertainty. The multidimensional cases were rerun at the same tightened settings, where thermal errors changed by at most 12% and frozen-volume disagreements by at most 0.11 percentage points. The practical lesson is that for this class of comparison the mesh is not the binding constraint and the iteration tolerance is. Where the VCN IIF pathway was added, additionally cases were rerun a third way (coupled model with both the nucleation pathways present).

#### Coupled model sequence of operations

An exact step by step description of the numerical iteration process is shown for one time step of a coupled-model simulation:

i. Start with the known temperature field, cell volumes, and PIF values from the end of the previous time step (call these the “old” values).
ii. Guess a trial new temperature field (initially, just reuse the old field as a first guess).
iii. Using this trial temperature, evaluate the material properties at each control volume (thermal conductivity and specific heat capacity as well as the current best estimate of the apparent heat capacity slope,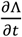.
iv. Assemble and solve the resulting (now linear, given the fixed apparent heat capacity from step iii) system for an updated trial temperature field (Thomas algorithm in 1D and one ADI directional sweep at a time in 2D/3D).
v. Using this updated trial temperature, run the cell-scale physics (described in Eqns. 2 to 8) forward by one time step, from the *old* cell state, to get a trial new Λ. Note that this involves solving the water-transport ODE and accumulating the nucleation-probability integrals for that one step.
vi. Compute the secant slope 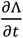 from the change in Λ (step v) over the change in temperature (step 4) just computed, and blend it with the previous iteration’s slope estimate using the under-relaxation or blending factor, ω, in the Picard iteration.
vii. Check whether Λ has stopped changing meaningfully compared to the previous iteration (the convergence test). If not converged, go back to step iii using the newly updated slope estimate (up to a maximum number of 200 iterations, after which the step is accepted regardless, as a practical safeguard).
viii. Once converged, accept the final trial temperature field, cell volumes, and PIF values as the “new” state, advance the simulation clock by Δ*t*, and move to the next time step (back to step i).

For the uncoupled (enthalpy) model, this same procedure applies, except step v is replaced by evaluating the equilibrium phase diagram (Eqn. 2 alone) directly at the trial temperature with no cell-scale physics involved. This makes the uncoupled model computationally lot less intensive and cheaper. From our computational experiments, roughly 15 times faster than the coupled model in this implementation.

#### Error metrics

The primary model disagreement is quantified by macroscale observables of the kind reported by Devireddy et al. [1], i.e., Δ*T*_*max*_ as the largest thermal-history difference over all probe locations and times. Other metrics included: (i) front-arrival error or the time at which the freezing front (the T = *T*_*ph*_ isotherm located by interpolation) first reaches half the depth, *d*/2 with error = (coupled time − uncoupled time) / uncoupled time × 100%. Front arrival error is negative when the coupled front arrives first, which it always did. Note that the relative front *position* was rejected as a useful metric because in thin samples the front crosses the domain almost immediately, so small absolute offsets at shallow depth read as large percentage errors. (ii) the final interface errors where the difference in the front position is determined as a percentage of the uncoupled position and was defined only if neither front has reached 97% of the domain. (iii) the peak osmotic lag or the largest value anywhere in the computational domain of the cell water retained, *x* minus the equilibrium value, Λ_*ec*_. (iv) the mean PIF in the frozen region or the average combined nucleation probability over the grid cells that are below *T*_*ph*_. (v) “changed by VCN” or cases where adding VCN changed Δ*T*_*max*_ by more than 0.05K and (vi) the frozen-volume gap (2D and 3D only) where at each of the recorded times is defined as the fraction of grid cells below and above *T*_*ph*_ in each model or the largest absolute difference between the two fractions. This is a clinically relevant quantity and, as shown in results, does not track the thermal error.

#### Numerical safeguards

Each trial temperature from the linear solve, and the intracellular freezing point 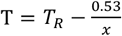are clipped to a floor of 1 K. Neither bound is reached by a converged solution and prevents a transient Picard iterate under an aggressive convective boundary from producing a non-physical temperature that would make the fractional-power nucleation exponent in Eqn. (7) undefined. Separately, the viscosity correlation of *η*(*T*) ∝ {(*T*/225) − 1}^−1.64^, is undefined below 225 K and we hold (*T*/225) −1 as 10^−6^ there, which suppresses SCN below −48 °C. Adding the two temperature floors left the reference results unchanged. The most demanding case (quenched 3D 1mm cube with both nucleation pathways active) required the tightening of the 2D/3D Picard settings from 20 iterations to 60 and tolerance from 10^−5^and 10^−6^. The other five multidimensional cases were then rerun at the tightened settings: frozen-volume disagreement changed by at most 0.001 and Δ*T*_*max*_ by −11.6% to +4.9%, so reported 2D/3D thermal errors carry roughly ±12% numerical uncertainty.

### SCALING ANALYSIS

As described so far, the two models (coupled and uncoupled) when computationally executed delineate, *after the fact*, how much they disagree for one specific case. What would be far more useful is a way to predict, *before* computing, roughly how big that disagreement will be, using only a handful of known tissue properties and the intended protocol. That is the goal of this section and is based on the physical intuition that the coupled and uncoupled models disagree because a real cell takes a finite amount of time to respond (by dehydrating) to a change in its surroundings, whereas the uncoupled model assumes that response is instantaneous. So the size of the disagreement should depend on a *competition* between two timescales: how long the cell actually needs to respond (denote this as *τ*_*osm*_ as derived below), and how long the tissue cell actually *has* to respond, given how fast the local temperature is falling (denote this as *τ*_*res*_, as defined below). If the cell has much more time than it needs, it keeps up, and the two models agree. If it has much less time than it needs, it falls behind, and the models disagree substantially. This scaling analysis combines these two timescales into a single non-dimensional grouping, ∏, and derives a closed-form formula to predict, *a priori*, how *large* the disagreement should be as a function of ∏. The closed-form formula is compared to the actual simulation predictions, as shown in the results section.

We start our scaling analysis by non-dimensionalizing the cell state as: 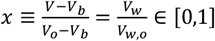 or 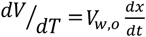. A solute parameter (*ε*) can also be defined as: 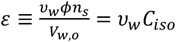, where *C*_*iso*_ is intracellular osmolality, *υ*_*w*_ is the partial molar volume of water, *ϕ* is the salt disassociation constant, *n*_*c*_ is the moles of salt in the cell. Numerically, s ≡ 5.4 × 10^−3^ ≪ 1, i.e., solute occupies a tiny fraction of the cell water. Thus, the logarithm *ln*[*x*/*x* + s] is a function of *x* alone. Setting the driving force in Eqn. (5) to zero defines *x*_*eq*_(*T*) as:

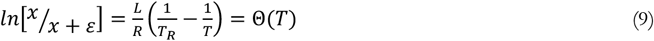

where *R* is the gas constant and *T*_*R*_ is the reference temperature (273.15 K). For modest sub cooling, Δ*T* = *T*_*R*_ − *T*, we can expand 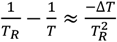 and 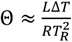. Since, 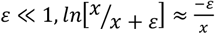. Therefore, 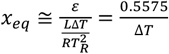, which essentially follows the Λ_*ec*_(*T*) shown in Eqn. (2). This result is not a coincidence and is a direct consequence of the fact that *at equilibrium the cell holds exactly the same unfrozen water fraction as the solution outside it*. Both are governed by the same colligative freezing-point depression and coupled and uncoupled models are identical in the quasi-static limit, and the entire discrepancy between them is solely due to a *kinetic lag*. Linearizing about the equilibrium: 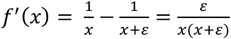 and *f*(*x*) −Θ ≈ *f*′(*x*)(*x* − *x*_*eq*_), such that:

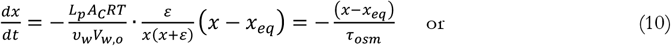

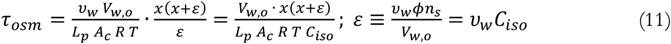

As *τ*_*osm*_ depends on state, we need a stated reference to define a number, by evaluating along the equilibrium locus at *x* = 0.5, which by Step 3 ties *T* to *x*: Δ*T* = 0.53/0.5 = 1.06 K and *T* = 272.09 K. For simplicity, the numerical model implements *τ*_*osm*_ at 272 K (rather than at 272.09K). This choice is justified by the fact that the latent heat release rate, *L* ∝ │*d*Λ/*dT*│ = 0.53/Δ*T*^2^, which is sharply peaked just below the freezing point, i.e., half the freezable water is gone by Δ*T* ≈ 1*K*.

The competing timescale is the local residence time and is defined as: 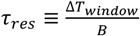, where Δ*T*_*window*_ is between *T*_*fo*_(272.62*K*) and 233.15 K (or the homogenous nucleation temperature of water) and *B* is the cooling rate. Note that thermal transport is not absent from the criterion, it is present through the cooling rate, *B* = *v*_*f*_ · *G* or front speed • local gradient. In the classical Stefan problem *v*_*f*_~ α/*x*_*f*_ (here α is the thermal diffusivity, m^−2^ · *s*^−1^) and *v*_*f*_~ (*T*_*fo*_ − *T*_*c*_)/*x*_*f*_, giving 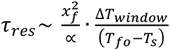, essentially a diffusion time built on the instantaneous front depth *x*_*f*_, not the domain size *L*_*d*_, and scaled by the fraction of the temperature drop spanned by the phase-change window. Early in the process *x*_*f*_ ≪ *L*_*d*_, and the two differ by orders of magnitude. A global alternative built on a domain-scale diffusion time, 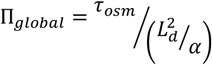 where *L*_*d*_ is the characteristic domain size/length, was tested and rejected (see results). Thus, the chosen final coupling group is ∏ = *τ*_*osm*_/*τ*_*rec*_. To recapitulate, ∏ ≪ 1 means the cell has far more time than it needs to keep up with the falling temperature and it stays close to equilibrium throughout, and the coupled and uncoupled models should agree closely. Conversely, ∏ ≫ 1 means the cell has less time than it needs and it falls badly behind, remaining mostly hydrated well past the point where equilibrium says it should have dehydrated, and the two models should disagree substantially.

Exactly the same style of analysis of evaluating the governing rate equation at a chosen reference state and determine where it predicts a transition, was applied to the nucleation Eqns. (5) to (8) and used to locate where each pathway (SCN or VCN) actually becomes fast enough to dominate over the other. We defined a *τ*_*SCN*_ ≡ 1/*SCN rate* and a *τ*_*VCN*_ ≡ 1/*VCN rate* as the characteristic time each pathway would need, at a given supercooling, to nucleate a cell, as detailed further in the results section.

#### Closed form lag law

We start this derivation by modeling the residence window as *x*_*eq*_(*t*) = 1 − *t*/*τ*_*re*s_ driving 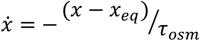, with *x*(0) = 1, the exact solution is: 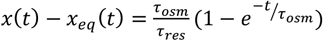. At the close of the window, *t* = *τ*_*res*_, with ∏ = *τ*_*osm*_/*τ*_*res*_ and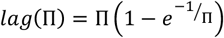. This gives:

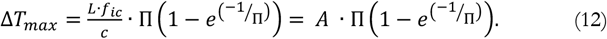

The limits are: lag → ∏ for ∏ ≪ 1 (error linear in ∏) and lag → 1 − 1/2 ∏ → 1 for ∏ ≫ 1 (saturation with all water retained). Admissible values of Π for a chosen error tolerance are obtained by inverting this relation numerically, as described later in the results section.

### INVERSE PROBLEM

To separate what a cell experiences from what the tissue does, one cell is integrated with Eqns. (5) to (8) forced by the uncoupled temperature history at mid-depth of a given protocol, carrying SCN and VCN as separate probabilities. We record the supercooling below the cell’s own *T*_*f*_ while it is still liquid (*PIF*^*tot*^ < 0.5), the temperature at which each pathway reaches 50%, and the unfrozen cell-water fraction, *x* at that moment. Supercooling is only meaningful before nucleation, so it is masked afterwards; and a pathway that fires after the cell has already frozen releases no further latent heat. For five protocols the coupled model was run with Ω_*o*_ or with baseline value of Ω_*SCN*_ multiplied by 0.01 to 100, and each run compared with the nominal one through the root-mean-square difference of the five probe histories, sampled every tenth step. This is the change an experiment would have to resolve, so it is quoted against thermocouple noise of 0.1 and 0.5 K. A “measurement” was, thus, produced by running the coupled model at the nominal Ω_*o*_ and *κ*_*o*_ for the 2 mm, 200 °C/min protocol and by adding an independent Gaussian noise of 0.1 K to every sampled probe reading. The two parameters were then scanned on a grid or 13 values of Ω/ Ω_*o*_ from 10^−3^ to 10^3^ crossed with 9 values of *κ*/ *κ*_*o*_ from 0.5 to 2, and a finer 7 × 7 grid over 0.5 to 2 and 0.90 to 1.10, with the “misfit” defined as the root-mean-square difference between each trial and the noisy measurement (117 and 49 coupled runs). Because misfit combines model separation and noise in quadrature, we report the separation 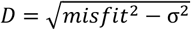, which is compared to the imposed noise. Note that no experimental data are involved in this inverse problem only synthetic computational data with added noise signals. Thus, this is a test of whether the parameters are recoverable. in principle and not a measurement of them (which is part of a companion study).

## RESULTS

The coupled and uncoupled solvers were first applied to the two cases (rat liver cooled at 5 °C/min and AT-1 tumor cooled at 50 °C/min) defined in Devireddy et al. [1]. The 1D solver was verified against the published results of Devireddy et al. [1] before any new computations were performed (Fig. 1). Specifically, Fig. 1(a) shows the freezing interface propagation in rat liver under the 5 °C/min “cryopreservation” protocol and Fig. 1(b) shows the thermal histories for rat liver at 0, 6, 16 and 32 mm. The coupled and uncoupled models differ by 1.4% in final interface position and are indistinguishable at this scale; the primary result of Devireddy et al. [1]. Quantities that follow from the tabulated properties without adjustment are reproduced as follows: maximum interface position 42.6 mm (reported ~40 mm) for cryopreservation and 21.8 mm (~21 mm) for cryosurgery; front onset at 128 s (~130 s); thermal histories terminating at −80, −70, −50 and −20 °C at 0, 6, 16 and 32 mm, matching the published values at every depth; and AT-1 tumor freezing 1.12% deeper than rat liver (~1.5%), an offset arising solely from the 210 versus 203 J/g difference in tissue latent heat. The central results of Devireddy et al. [1] were, thus, recovered with the coupled and uncoupled models differing by 0.65 to 1.35% across the four published cases, within the reported 3% bound. Convergence within a model was established by halving the time step (0.1 → 0.05 s, changing the coupled–uncoupled difference from 8.74% to 8.59%) and by increasing the Picard cap (100 → 300 iterations, 8.78% → 8.75%). The 2D/3D solver reproduces the 1D solver to ~15% when run in a quasi-1D configuration, which is adequate for regime mapping (their primary function in this study) but not for quantitative front prediction. The reimplementation was written from the published equations and tables alone, in Python, without reference to the original Fortran code, which is no longer readily executable. The agreement reported above is therefore an independent check on the published description of the model, as well.

**Fig. 1:**
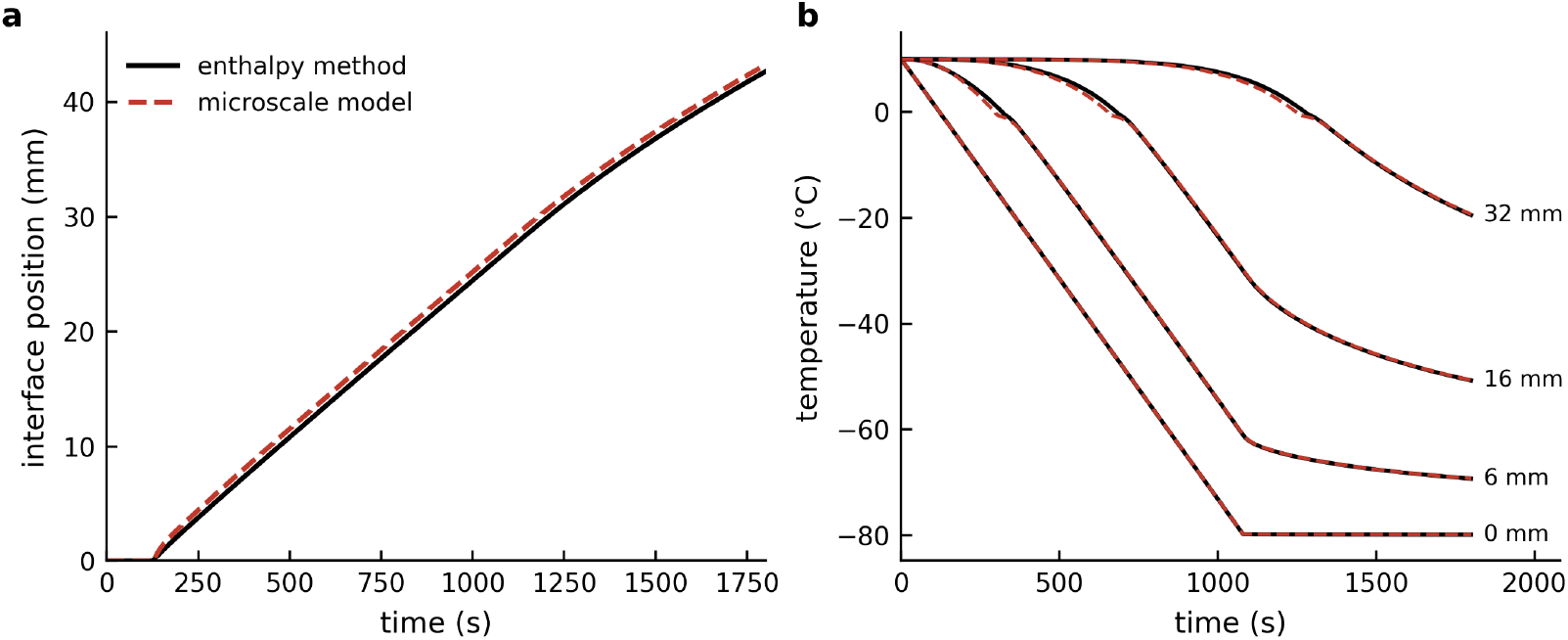
A comparison of the results from Devireddy et al. [1] with the results from the code developed, independently in the present study. The simulation domain is an 8 cm slab of rat liver under the “cryopreservation” protocol of Devireddy et al. [1], i.e., cooled on one face at 5 °C/min to a hold at −80 °C. **Fig. 1(a)** shows the position of the freezing front against time. The solid line is the uncoupled enthalpy model and the dashed line is the coupled model. Both the models agree to within ±3% and with the earlier results. **Fig. 1(b)** shows the temperature against time at four depths (0, 6, 16 and 32mm). Solid lines are the uncoupled model and dashed lines the coupled model. The front reaches ~42.6 mm in the current study against ~41.7 mm in the original study, and begins to advance at ~128 s against the earlier study value of ~130 s. The two computational models (this study and the earlier study) differ by ~1% in final front position and are statistically equivalent.

Fig. 2 shows the origin of the osmotic lag and the region where it is prevalent. Specifically, Fig. 2(a) compares the intracellular equilibrium obtained by solving the transport equation for zero driving force, *x*_*eq*_ = 0.5575/Δ*T*, with the extracellular phase diagram, Λ_*ec*_ = 0.53/Δ*T*. Not unexpectedly, the two curves agree to 5.2% over the range 0.6 to 12 K of sub-cooling. The coupled and uncoupled models are therefore identical in the quasi-static limit, and every difference between them is a kinetic lag and nothing else. Fig. 2(b) shows unfrozen-fraction profiles at *t* = 400 secs in the 5 °C/min rat liver tissue cryopreservation case. Behind the freezing front the retained cell water tracks the equilibrium curve closely. However, within approximately 1.5 mms of the front (the width over which the lag exceeds half its peak value) the two diverge, reaching a maximum difference of 0.40 at 7.95 mm, where cells retain 84% of their water while the equilibrium value is 44%. The IIF probability is zero throughout this band. The relative disagreement in the freezing front position in this same case falls from +8.9% at *t* = 400 secs to +1.4% at *t* = 1800 secs. If the SCN IIF parameters were to “nucleate” ice in this regime the coupled and uncoupled thermal profiles would have diverged in the earlier study of Devireddy et al. [1]. The lag is not distributed through the frozen region, instead it is prevalent in a narrow fixed band pinned to the freezing front. This leads to a phenomenon where in the earlier 8cm domains of Devireddy et al. [1] the front travels 43 mm and the 1.5 mm lag zone band is 3.5% of it, and the models agree to 1.35%. However, in a 1 mm sample the band is comparable to the entire domain, so there is no bulk of equilibrated frozen tissue to average it away, and the same physics produces errors of tens of percent. Thus, the geometry *dilutes* the discrepancy where, somewhat paradoxically, the smaller domain sizes reveal what the larger domain scales conceal. A scaling of *τ*_*osm*_ with *x*(*x* + ε) demonstrates a second effect i.e., a fully hydrated cell at the freezing front is roughly an order of magnitude slower to respond than a partly dehydrated one behind it (data not shown). There is also upturn in the lag at small *x* and is a second effect worth noting. This is due to the Arrhenius dependence of *L*_*p*_ (*E*_*Lp*_ = 265 kJ/mol for rat liver) eventually freezing the membrane shut faster than the shrinking volume speeds relaxation.

**Fig. 2:**
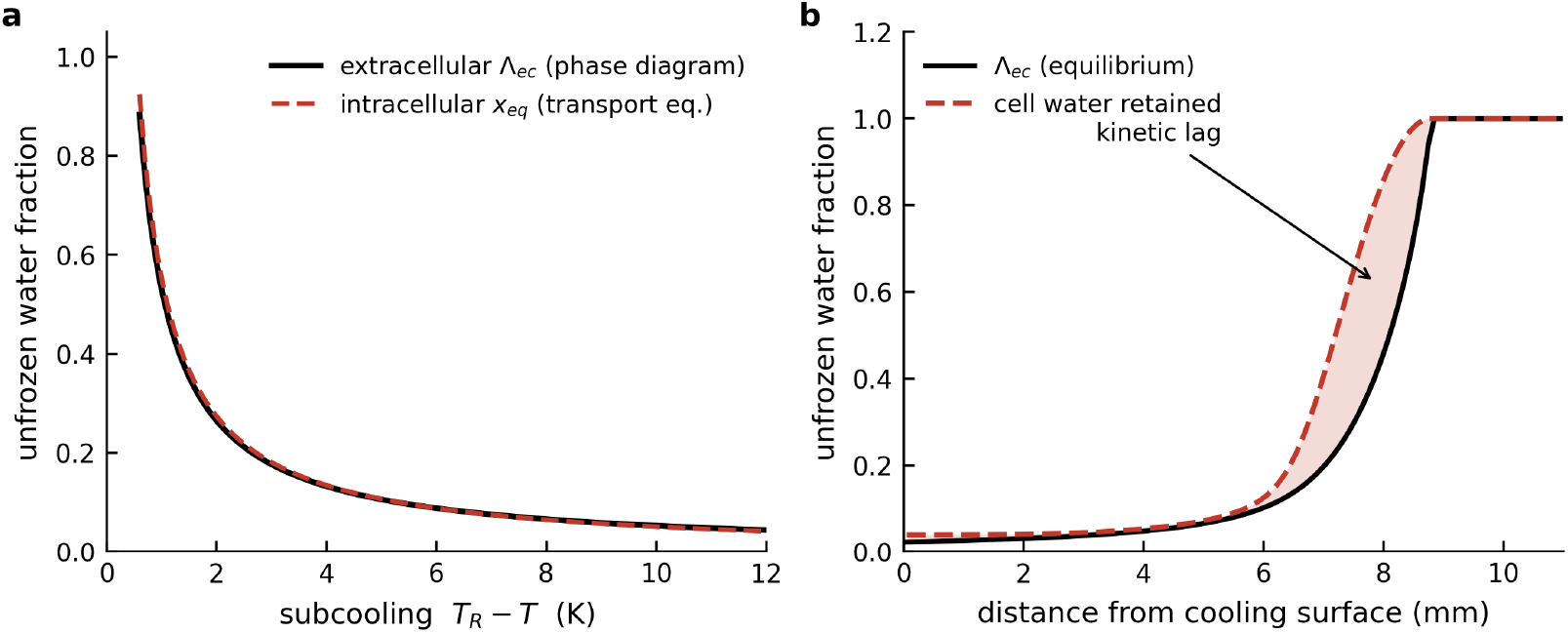
The figure demonstrates the fundamental underlying cause for the possible difference between the coupled and uncoupled models. **Fig. 2(a)** shows the equilibrium cooling line, i.e., the unfrozen water fraction a cell would hold if its transport equation were solved to equilibrium at each temperature. The dashed line is the extracellular phase diagram used by the uncoupled model. The two agree to ±5.2% between 0.6 and 12 K of sub-cooling, so the two models share the same thermodynamics and differ only in timing. **Fig. 2(b)** shows a snapshot at 400 s during the “cryopreservation” protocol, i.e., cooled on one face at 5 °C/min to a hold at −80 °C. The dashed line is the equilibrium unfrozen fraction, the solid line is the water actually retained by the cells, and the shaded area between them is the kinetic lag. The lag reaches 0.40 at a depth of 7.95 mm, where cells still hold 84 per cent of their water while equilibrium calls for 44 per cent. The intracellular ice probability is zero across this band, so the retained water is trapped rather than frozen in place. The band is about 1.5 mm wide, set by how far the front advances while one cell relaxes, and it does not grow as the sample or the run gets longer. This fixed lag band width is why a large domain hides a discrepancy that a small one exposes.

Fig. 3 shows the efficacy of the ∏ = *τ*_*osm*_/*τ*_*rec*_ (Fig. 3a) over the competing choice of 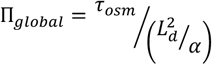(Fig. 3b) for the 51 protocol cases in which sample size and cooling severity both vary. The error rises monotonically across four decades of ∏, with a Spearman rank correlation, *S* = 0.92 (Fig. 3a) while the same data plotted against ∏ _*gIobaI*_ gives a Spearman rank correlation, *S* = 0.24 (signifying a model agreement that is barely better than no relationship at all). The two cases highlighted in Fig. 3(b) share an identical ∏ _*g*I*oba*I_ = 0.85, i.e., the same 2 mm sample and diffusivity, differing only in programmed cooling rate but produce errors of 0.22 and 9.45 K (~43x difference). A domain-scale diffusion time cannot resolve the results as it contains no cooling rate dependence, and cooling rate is the dominant control. Another reason is locality, i.e., the osmotic competition is settled inside one ~20 μm Krogh cylinder during the seconds while the front sweeps past, and whether tissue extends 2 mm or 8 cm beyond that point is information the cell has no access to and no dependence upon. Note that the thermal transport is not absent from the ∏ criterion and it enters through *τ*_*res*_ via the local cooling rate locally, and not globally, as shown earlier.

**Fig. 3:**
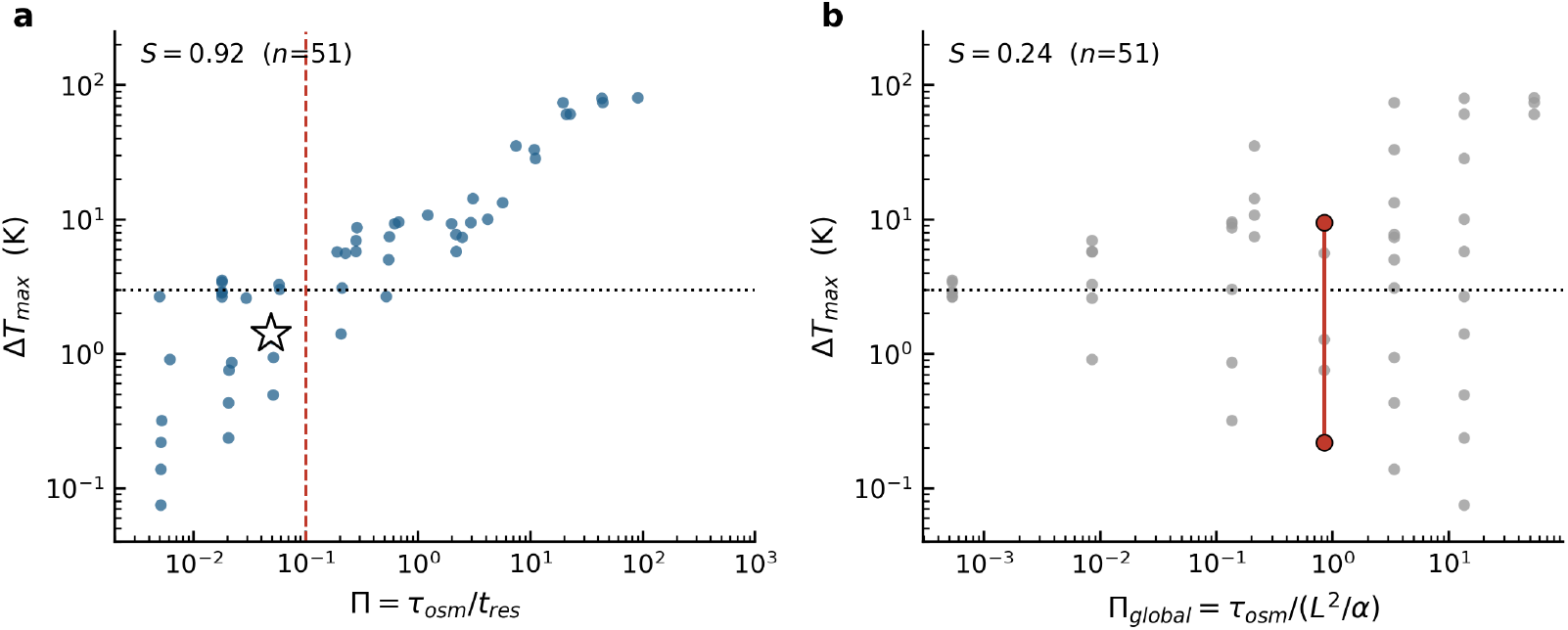
The figure shows the scaling of the data against two possible ∏ groups, i.e., **∏** = *τ*_*osm*_/*τ*_*rec*_ or 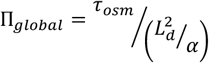, as described in the text. Both figures show the **∏** value on the x-axis and the corresponding model calculated, ΔT_*max*_. **Fig. 3(a)** plots ΔT_*max*_ vs. ∏ = *τ*_*osm*_/*τ*_*rec*_ with a Spearman rank correlation (*S*) value of 0.92, suggesting a very good correlation between both the variables. The dashed vertical line marks ∏=0.1 and the dotted horizontal line marks the 3 K bound adopted in this study. The open star is the operating point of Devireddy et al. (2002), at Π = 0.049 and 1.41 K. **Fig. 3(b)** orders the same data against a group built instead on domain-scale thermal diffusion 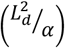. The two red points shown on the graph have the same 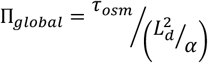 value of 0.85, yet the ΔT_*max*_ values vary by a factor of ~40 (0.22 K to 9.45 K). The two red data points differ only in the imposed cooling rate. This lack of correlation between the two is demonstrated by the very low Spearman rank correlation (*S*) value of 0.24.

Table 3 summarizes all the 124 one-dimensional cases classified by ∏. Median Δ*T*_*max*_ rises from 0.88K for ∏ < 0.03 to 33K for ∏ > 10, and median front-arrival error from −1.3% to −41.3%. The median peak osmotic lag rises correspondingly from 0.02 to 1.00, saturating at complete water retention. A 3K bound on Δ*T*_*max*_, adopted in this study as a thermal-history analogue of the 3% interface criterion of the earlier study [1] and a threshold not stated in that work, is crossed by the bin medians between the ∏ = 0.03 *to* 0.1 (median 1.27 K) and the ∏ = 0.1 *to* 0.3 (median 4.9 K). As stated earlier, the operating point of Devireddy et al. [1] lies at Π = 0.049 with Δ*T*_*max*_ = 1.41 K (and shown as a * in Fig. 3a). The bin medians conceal three exceptions, i.e., the controlled-rate cooling of the 80 mm slab at 20, 50 and 200 °C/min which gives Δ*T*_*max*_ = 2.7, 3.4 and 3.5 K, respectively all at ∏ = 0.018 and all above the 3K bound despite lying in the lowest bin. All three cases share an identical ∏ across a tenfold range of cooling rate, which shows that the median residence time defining ∏ is set here by deep, diffusion-controlled probe locations, whereas Δ*T*_*max*_ arises near the surface, where cooling is fastest and the local ratio is much larger. However, their front-arrival errors remain small (1.8 to 2.0%). As described in the earlier study, the coupled front arrives earlier than the uncoupled front, in every case investigated.

**Table 3:**
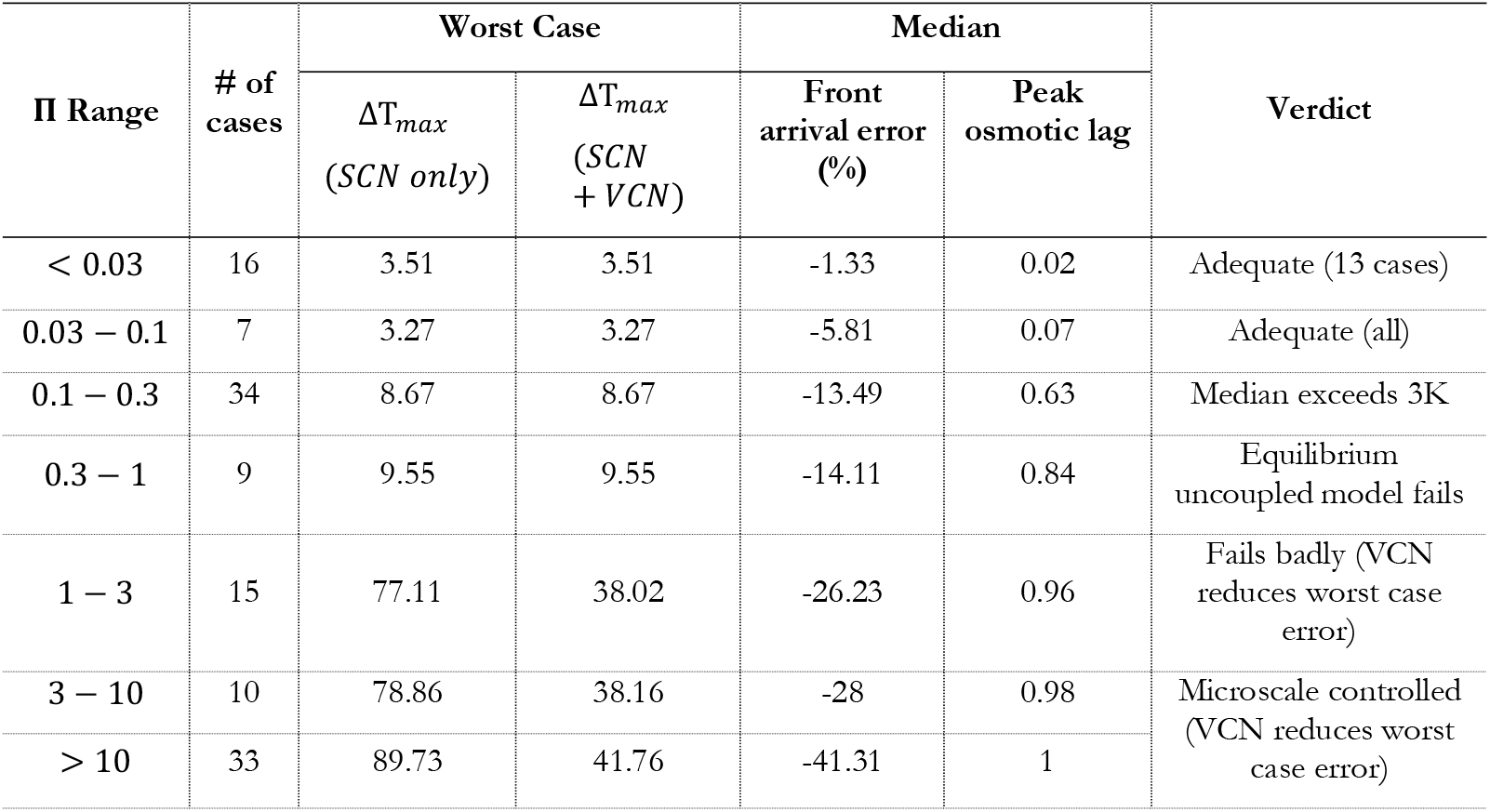
The 124 one-dimensional cases binned by the coupling group. Values are converged (Picard tolerance 10^−8^ with 200 iterations). The front-arrival error is negative because the coupled front always arrives first.

| $\Pi$ Range | # of cases | Worst Case | | Median | | Verdict |
| --- | --- | --- | --- | --- | --- | --- |
| | | $\Delta T_{max}$<br>(SCN only) | $\Delta T_{max}$<br>(SCN + VCN) | Front arrival error (%) | Peak osmotic lag | |
| < 0.03 | 16 | 3.51 | 3.51 | -1.33 | 0.02 | Adequate (13 cases) |
| 0.03 – 0.1 | 7 | 3.27 | 3.27 | -5.81 | 0.07 | Adequate (all) |
| 0.1 – 0.3 | 34 | 8.67 | 8.67 | -13.49 | 0.63 | Median exceeds 3K |
| 0.3 – 1 | 9 | 9.55 | 9.55 | -14.11 | 0.84 | Equilibrium uncoupled model fails |
| 1 – 3 | 15 | 77.11 | 38.02 | -26.23 | 0.96 | Fails badly (VCN reduces worst case error) |
| 3 – 10 | 10 | 78.86 | 38.16 | -28 | 0.98 | Microscale controlled (VCN reduces worst case error) |
| > 10 | 33 | 89.73 | 41.76 | -41.31 | 1 |  |

Fig. 4(a) shows *τ*_*SCN*_ and *τ*_*VCN*_ as a function of supercooling below the intracellular freezing point. Evaluating these directly from the Eqns. (5) to (8), using the published rat liver parameters, at the same reference state (*x* = 0.5) used for *τ*_*osm*_, the value for *τ*_*SCN*_ falls from around 2 × 10^12^ secs at 2 degrees of supercooling to 2.4 secs at 10 degrees of supercooling, an enormous change over a small degree of supercooling change. In contrast, *τ*_*VCN*_ remains above 10^8^ seconds until roughly 30 degrees of supercooling, and only drops to ~1 sec timescales beyond about 38 degrees of supercooling (or around 233K). This suggests, as postulated elsewhere, that the two ice nucleation pathways occupy different, non-overlapping supercooling ranges. Note that the VCN rate peaks 110 degrees of supercooling (or −110 °C) and falls again at colder temperatures, due to the *T*^3^ factor in the denominator of the exponent in Eqn. (8) outweighing the growing supercooling drive. This observation is an established feature of classical nucleation theory [78]. For SCN the picture is different, and partly artificially due to holding the viscosity constant at ~10^6^ Pa · s for *T* < 225K (as stated earlier, to avoid numerical anomalies). This switches SCN off below 225K and comes from the clipping the empirical fit of viscosity outside its range and not from nucleation physics, *per se*. However, it is not too much of a stretch to assume that viscosity below 225K is a very high value and is probably in the range of 10^6^ Pa · s [47].

**Fig. 4:**
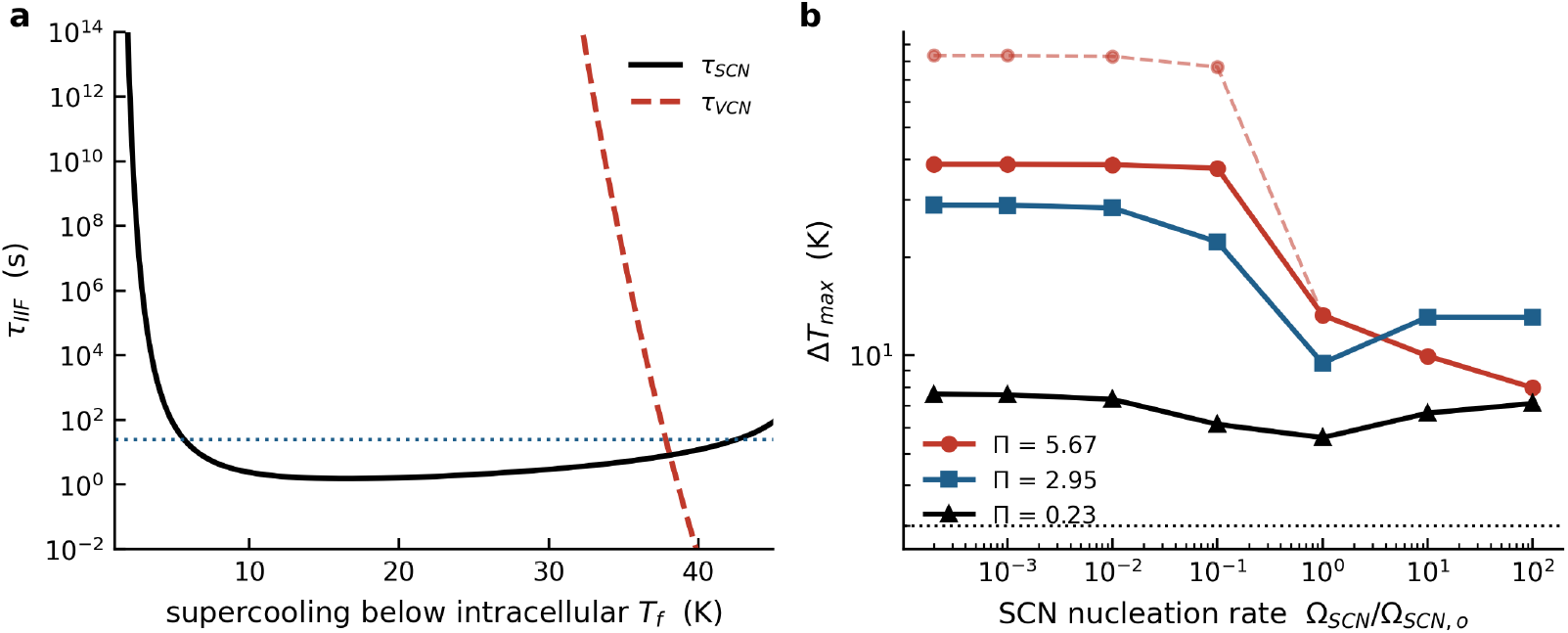
The figure shows the disparate temperature regions within which the two IIF phenomena (SCN and VCN) are manifested as well as the consequence non-overlapping behavior on the latent heat release during freezing. **Fig. 4(a)** shows the characteristic nucleation time (y-axis) against the degree of supercooling below the cell’s own freezing point (on x-axis), evaluated at half dehydration; see text for further details. The solid line is the SCN pathway while the dashed line is the VCN pathway. The SCN time falls from 2 × 10^12^ s at 2 K of supercooling to ~2.4 s at 10 K of supercooling, an extraordinary drop in a narrow supercooling window. The VCN time stays above > 10^8^ s until about 30 K of supercooling and only becomes “active” beyond ~38 K, so the two occupy distinct and separate temperature ranges and they almost act as distinct “on-off” switches. The dotted horizontal line marks the osmotic relaxation time (*τ*_*osm*_) of 24 s for rat liver tissue and the timescale to which these two values are competing against. Note the process with a lower relaxation time value dominates the freezing process. **Fig. 4(b)** shows the maximum thermal difference against the surface nucleation rate, expressed as a multiple of its measured value, at three fixed values of the ∏ coupling group. Solid lines include both SCN and VCN nucleation pathways while dashed lines include the SCN pathway only. The dotted horizontal line is the 3 K bound. Note that intracellular water that fails to leave the cell can still freeze in place, which releases latent heat that the equilibrium model assumes was already expended. As expected, suppressing ice nucleation therefore produces the largest errors and accelerating it restores agreement between the equilibrium (uncoupled) and the micro-scale (coupled) models. The dashed and solid curves converge once the SCN rate alone is fast enough to saturate nucleation or release all of the intracellular latent heat, which is why the second VCN pathway matters only where the first (SCN) is weak or suppressed.

Adding VCN changes the reported error in 24 of the 124 one-dimensional cases and in 2 of the 6 multidimensional cases; the remaining 100 and 4 cases, respectively, are unchanged to within 0.05 K. Every affected case uses a convective liquid-nitrogen quench while the controlled-rate-freezer protocols are unaffected at any cooling rate tested (0.5– 200 °C/min), including cases reaching ∏ > 100 through enlarged cell size. The affected cases fall into two groups: (i) where SCN nucleation is otherwise weak and the reduction is still significant. For example at ∏ = 10.8, with SCN suppressed entirely, VCN alone still reduces the error from 89.7 K to 41.8 K while at nominal SCN strength the same case falls from 33K to 29.3K. (ii) where SCN alone is already strong, i.e., larger cells, whose greater membrane area raises the SCN rate directly and where VCN adds nothing further. In the cell-size sweep at severe cooling, increasing linear size from 0.5× to 10× reduces Δ*T*_*max*_ at the reference permeability from 78.9 to 9.3K under SCN alone. The same cases with VCN alone show Δ*T*_*max*_ ranging from 38.2 to 9.3K. Here, the largest cells are unchanged, while the smallest (whose small membrane area makes SCN weakest) gain most from the volume-driven pathway, so VCN compresses the size dependence rather than leaving it intact. Fig. 4(b) shows this directly: at fixed ∏, SCN-only (dashed) and SCN+VCN (solid) curves converge once the SCN rate is turned up enough to saturate nucleation on its own. The frozen-volume-fraction disagreement is unaffected by the inclusion of VCN in every one of the 6 multidimensional cases, even where the thermal error falls substantially. VCN closes part of the thermal-history gap without changing the predicted extent of the frozen region. Therefore, the formation of intracellular ice releases the latent heat that osmotic efflux failed to deliver, and thereby rescues the equilibrium model. The failure condition is therefore a window requiring two conditions together (slow dehydration and slow nucleation) not a threshold in either variable alone. The two routes are not independent alternatives but causally linked as a cell that dehydrates successfully keeps tracking the local temperature and never accumulates supercooling, whereas a cell that fails to dehydrate holds the intracellular water and it’s phase-change temperature stays pinned near −0.5 °C while the surrounding temperature falls. Osmotic failure is precisely what generates the supercooling that triggers nucleation, which is why the error saturates rather than growing without bound. The practical implication is counter-intuitive and suggests that a protocol optimized to suppress IIF for cell survival simultaneously invalidates the heat-transfer model used to design it.

Fig. 5(a) shows frozen volume fraction against time for an 8 mm cube cooled on all six faces at 20 °C/min (∏ = 0.22). The models disagree by 25 percentage points at peak. Cooling on six faces rather than one superposes heat extraction and shortens the local residence time relative to the 1D equivalent at the same nominal cooling rate, so dimensionality is not a numerical refinement but part of the regime criterion or identical tissue under an identical programmed rate can sit on either side of the boundary depending solely only on how many faces are cooled. Fig. 5(b) reports the peak frozen-volume disagreement for all the six multidimensional cases, ordered by ∏ ranging from 1.8 percentage points for the axisymmetric cryoprobe (∏ = 0.33), 25.4 for the 8 mm cube, 32.8% for the 4 mm cube (∏ = 0.66), 38.3 for the block corner (∏ = 0.67), 39.6 for the vitrification carrier (∏ = 97) and 31.7 for the 1 mm cube (∏ = 131). These figures are unchanged by the inclusion of VCN. As expected, the thermal and volumetric measures do not correspond. The 2D cryoprobe gives Δ*T*_*max*_ = 4.7 K with a 2-point volume gap because radial divergence spreads the lag band over an expanding front. In contrast, the 2D block corner gives a comparable 7.4 K but with a 38-point gap as the corner concentrates it. Thus, a model can match thermal histories to a few degrees while misplacing the frozen boundary badly. One qualification applies to the 2D cryoprobe, i.e., the probe spans the full height of the domain and only its surface is cooled, so the solution is uniform along the axis (to 10^−11^K) and the case is effectively one-dimensional radial problem and it only tests the radial divergence, not tip effects. All six cases exceed the 3K bound on thermal error although with VCN included, the two convective-quench cases (vitrification carrier and 1 mm cube) decrease the Δ*T*_*max*_ from 71.3 to 34.6 K and from 70.8 to 44.2 K respectively. The other four are unchanged, consistent with the protocol-type pattern described earlier.

**Fig. 5.**
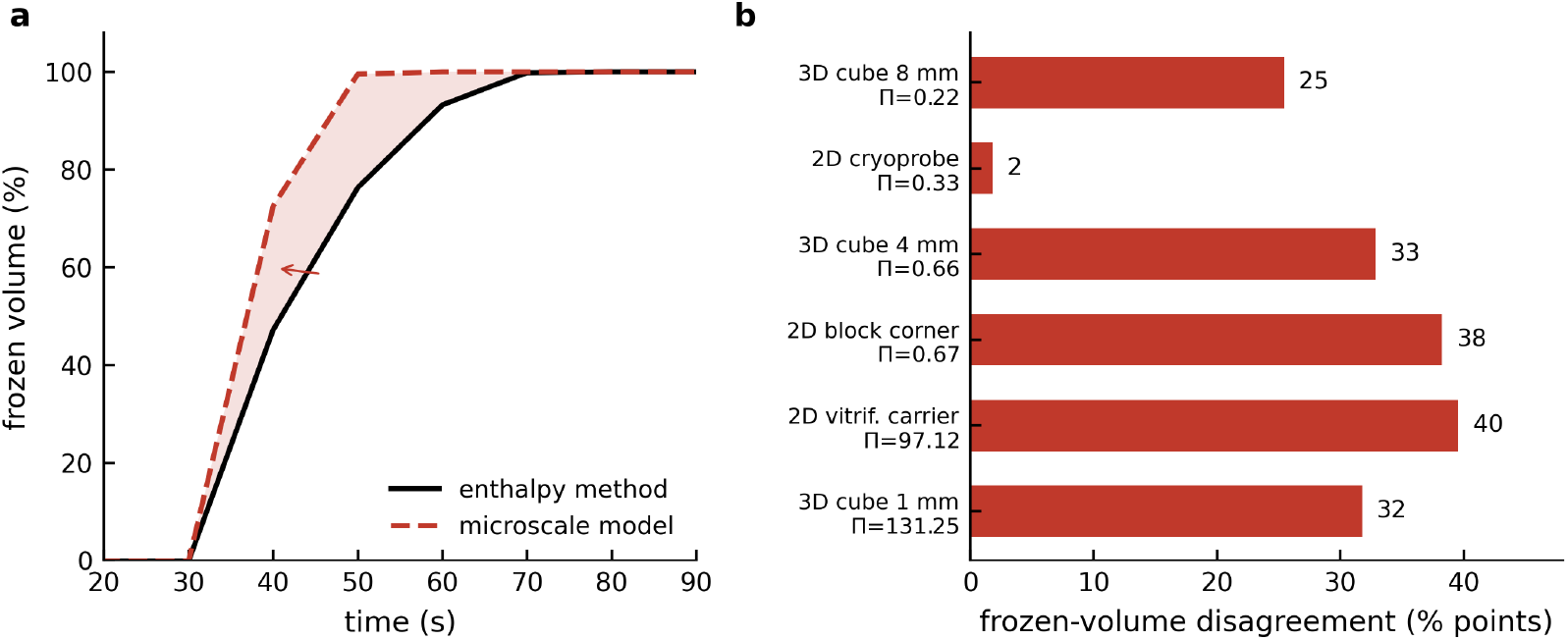
This figure shows the consequences of expanding the models to higher dimensions and the disagreement in the frozen volume fraction. **Fig. 5(a)** shows the % frozen volume fraction against time in seconds for an 8 mm cube cooled on all six faces at 20 °C/minute, computed as an octant by symmetry on a 40 × 40 × 40 grid. The solid line is the uncoupled (equilibrium cooling) model and the dashed line is the coupled (microscale) model. The arrow marks the largest disagreement of 25.4 percentage points. Cooling on six faces rather than one superposes heat extraction and shortens the local residence time (*τ*_*res*_) without changing any material property, so a cooling protocol can cross the boundary between the uncoupled and coupled models through geometry alone. **Fig. 5(b)** lists the peak frozen-volume disagreement (in % points) between both the models for all the two- and three-dimensional cases explored in the current study. The 2D and 3D cases are ordered by the coupling group, with the exact percentage point difference value printed beside each bar. Note that the disagreement is large in a quantity that matters for cryosurgical planning and one that is invisible in a one-dimensional thermal history. As stated, in the text the thermal-time histories and volumetric measures do not track each other.

Fig. 6(a) plots all one-dimensional results against the lag law, derived earlier as: 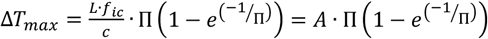. With SCN only, least-squares fits give *A* = 29.4 K at nominal nucleation rate and *F* = 69.1 K with nucleation suppressed entirely (root-mean-square residual 17.5 K over 124 points). With SCN and VCN combined, the same fit gives *A* = 20.3 K (a lower ceiling) and a smaller residual, 8.1 K over the same 124 points. This suggests that including the second VCN pathway does not merely reduce the worst case, it makes the single-parameter law describe the data more precisely. The largest error observed in any simulation is 89.7 K under SCN alone and 41.8 K with both SCN and VCN pathways. The predicted ceiling from the intracellular latent-heat content, 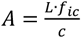 is 94K against an observed saturation near 90K. This acts as an independent confirmation that the mechanism was correctly identified as the number was never fitted (nor used) in our simulations. Fig. 6(b) gives the relation inverted for admissible ∏ against accepted error, tabulated in Table 4 for all three cases. For a 3K tolerance the lag law returns ∏ _*to*I_ = 0.148 with both SCN and VCN pathways active, a value of 0.102 with SCN alone, and a value of 0.043 with neither being active. Corresponding number for a 1K tolerance are 0.049, 0.034 and 0.014, respectively. While for a 10K tolerance the value are 0.613, 0.363 and 0.145, respectively. Intriguingly, the inclusion of the VCN relaxes the design criteria as the additional nucleation pathway provides real physical relief for the unfrozen fraction that the SCN-only analysis could not access. Inverting 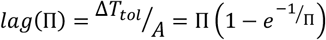 turns the result into a design tool, i.e., choose a tolerance, read off ∏ _*to*I_, and compare against ∏ computed from tabulated cell properties (*τ*_*osm*_) and the intended protocol (*τ*_*res*_), before committing to a coupled simulation costing roughly 15× more.

**Table 4:**
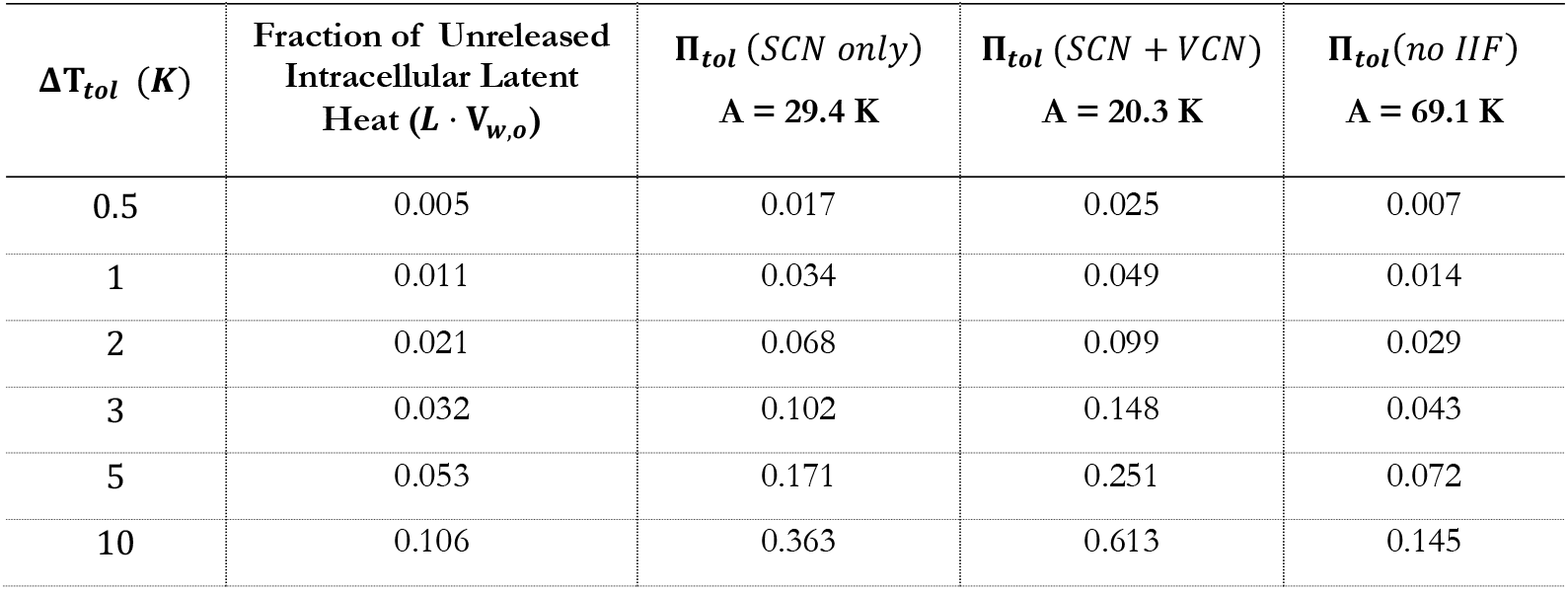
Admissible coupling group for a chosen error tolerance, obtained by inverting the lag law. The three columns correspond to three assumptions about ice nucleation relief.

| $\Delta T_{tol}$ (K) | Fraction of Unreleased Intracellular Latent Heat ($L \cdot V_{w,o}$) | $\Pi_{tol}$ (SCN only)<br>A = 29.4 K | $\Pi_{tol}$ (SCN + VCN)<br>A = 20.3 K | $\Pi_{tol}(no IIF)$<br>A = 69.1 K |
| --- | --- | --- | --- | --- |
| 0.5 | 0.005 | 0.017 | 0.025 | 0.007 |
| 1 | 0.011 | 0.034 | 0.049 | 0.014 |
| 2 | 0.021 | 0.068 | 0.099 | 0.029 |
| 3 | 0.032 | 0.102 | 0.148 | 0.043 |
| 5 | 0.053 | 0.171 | 0.251 | 0.072 |
| 10 | 0.106 | 0.363 | 0.613 | 0.145 |

**Fig. 6.**
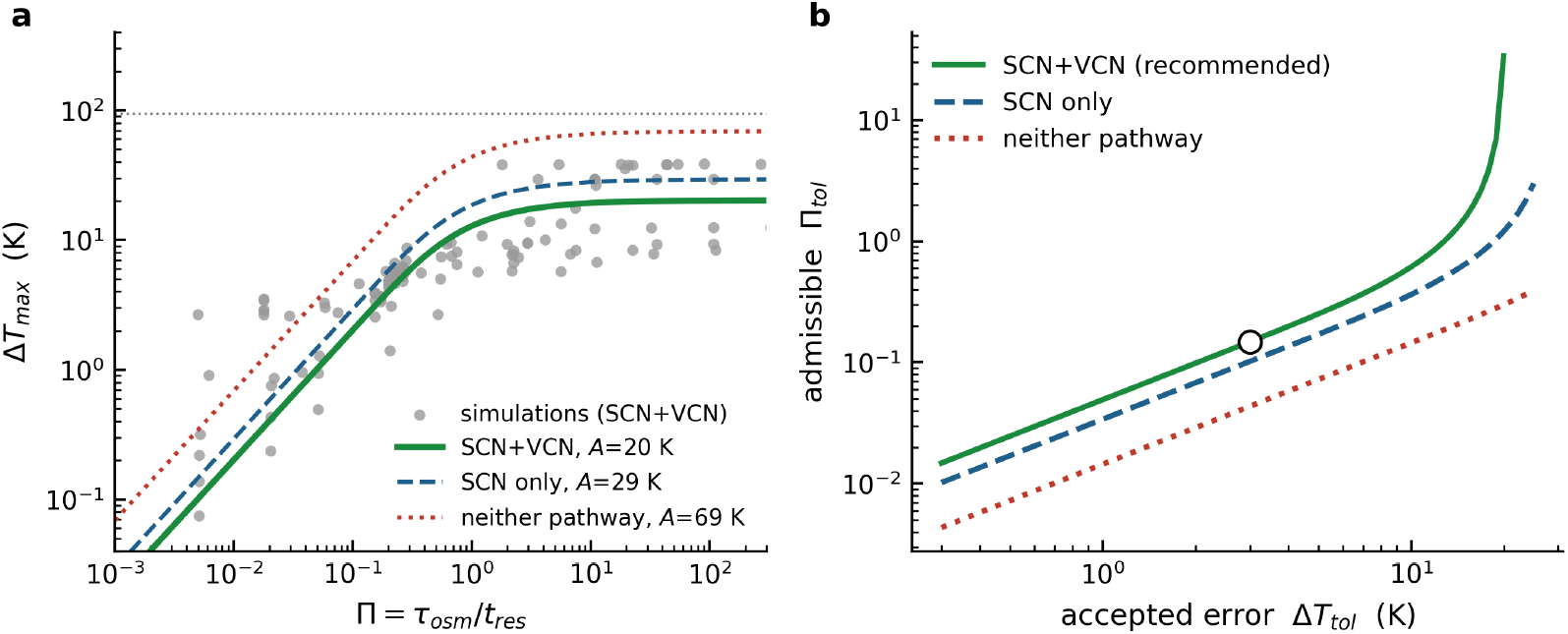
This figure demonstrates the utility and use of the analytical lag law derived in the text. **Fig. 6(a)** plots all the one-dimensional ΔT_*max*_ results against the lag law, with the coupling group (∏) on the horizontal axis. Each data point is a simulation. The three curves are the law fitted with nucleation pathways both (SCN and VCN) active, with the surface pathway (SCN) only, and with neither being active, giving ceilings of 20.3, 29.4 and 69.1 K. The dotted horizontal line is the ceiling predicted from the intracellular latent heat alone, 94 K, which was never fitted. The predicted 94 K ceiling (calculated using tissue properties and one that was never fitted) agrees quite closely with the observed saturation near 90 K. This agreement provides independent support and validation to the derived analytical lag law mechanism. Note that including the second nucleation pathway (SCN) lowers the ceiling and also improves the fit, with the residual falling from 17.5 K to 8.1 K, which further supports our assertion that the lag law captures the underlying mechanism rather than absorbing scatter. **Fig. 6(b)** shows the lag law relation inverted, giving the largest admissible coupling group (∏) for a chosen error tolerance (ΔT_*to*I_), for the same three assumptions of IIF nucleation. Essentially, panel (b) acts as a design tool. The dot in the panel represents a ΔT_*to*I_ of 3 K with the admissible ∏ group being read-off the y-axis as 0.148 with both SCN and VCN pathways being active. The other admissible ∏ numbers are 0.102 with just the SCN pathway being active and a value of 0.043 with neither pathways being active. If the calculated value of ∏ = *τ*_*osm*_/*τ*_*rec*_ is below this admissible ∏ value then the design engineer can resort to the use of equilibrium (uncoupled) freezing model while the opposite condition will necessitate the use of the coupled micro-macroscale freezing model; see text for further details.

Specifically, using the design choice tool takes four steps. First, compute *τ*_*osm*_ from tabulated cell properties using the expression Eqn. (11), which for rat liver gives 24 s. Second, obtain *τ*_*res*_ for the intended protocol, either from the protocol itself or from a single inexpensive run of the uncoupled model, and form ∏ = *τ*_*osm*_/*τ*_*rec*_. Third, decide what error is tolerable (Δ*T*_*to*I_) for the purpose at hand. Fourth, read ∏ _*to*I_ for the predetermined (Δ*T*_*to*I_) from Fig. 6(b) or Table 4. If ∏ is below ∏ _*to*I_, the equilibrium model is adequate and the coupled model is not worth its cost. If the read ∏ from Fig. 6(b) is above ∏ _*to*I_,, the coupled model is required to fully capture the underlying physics. Two worked examples further illustrate the use of our design tool. The cryopreservation case of the original study by Devireddy et al. [1] has ∏ = 0.049 for rat liver and 0.044 for AT-1 tumor. At a 3K tolerance or (Δ*T*_*to*I_ = 3*K*) the corresponding ∏ _*to*I_ value from Fig. 6(b) is 0.148. So the calculated ∏ values are comfortably below the limit and the equilibrium model suffices, which is what the original study found by direct comparison and what the simulation confirms with an error of 1.4 K. The 8 mm cube cooled on six faces at 20 °C/min has ∏ = 0.22, which exceeds the ∏ _*to*I_ value of 0.148 and so the criterion calls for the coupled model. The simulated error is 6.9 K, which is indeed above tolerance.A 3K tolerance returns ∏ _*to*I_ = 0.102 (SCN alone) or 0.148 (SCN + VCN) and both these values fall inside the interval bracketed by the simulations. The closed form law and the simulations therefore agree without any parameter tuning, although the binning locates the boundary only to within a factor of three. This last result strongly supports (if not completely vindicate) our analytic law, as the simulations and the derived law both agree without any additional parameter tuning. Therefore, for most tissues, where both nucleation pathways are available, the SCN plus VCN curve is the most appropriate choice. The curve with neither pathway active should be used when the nucleation behavior of a tissue is unknown, or when a protocol has been designed specifically to suppress intracellular ice, since such a protocol places the user on that curve by construction

The conditions under which the micro-scale freezing processes (water transport and the SCN/VCN IIF) modify the macro-scale freezing response is dependent on the underlying governing parameters (*L*_*pg*_, *E*_*Lp*_, Ω_*SCN*_, *κ*_*SCN*_, Ω_*VCN*_ and *κ*_*VCN*_). Although the water transport parameters (*L*_*pg*_, *E*_*Lp*_) can be determined for tissue cells using a combination of low temperature microscopy and calorimetric methods [34,36–43], the corresponding SCN and the VCN parameters are, as yet, unknown a no experimental techniques exist to determine them. That asymmetry inspired us to explore an inverse approach, where we synthetically or computationally impose a known cooling history, measure temperature, and ask which nucleation parameters reproduce it. Three calculations are presented to test whether this is possible in principle. Fig. 7a shows that changing, Ω_*o*_ by a factor of ten alters the predicted thermal history by 0.29 to 0.42 K at 20 °C/min, 0.54 to 1.02 K at 60 °C/min and 0.84 to 3.45 K at 200 °C/min, against an anticipated thermocouple noise of order 0.1 to 0.5 K. At 2°C/min the same change moves the history by 0.01K and is unmeasurable as the cell is ~ 97% dehydrated when it nucleates, so freezing it releases almost no latent heat to measure and analyze. The signal is also asymmetric, being larger when, Ω_*o*_is under-estimated than over-estimated, because nucleation that is already effectively instantaneous cannot be made to proceed much faster. Fig. 7(b) shows the cooling rate required for a 2mm slab for the measured temperature signal to be above the thermocouple noise or cooling rates above 10 °C/min.

**Fig. 7.**
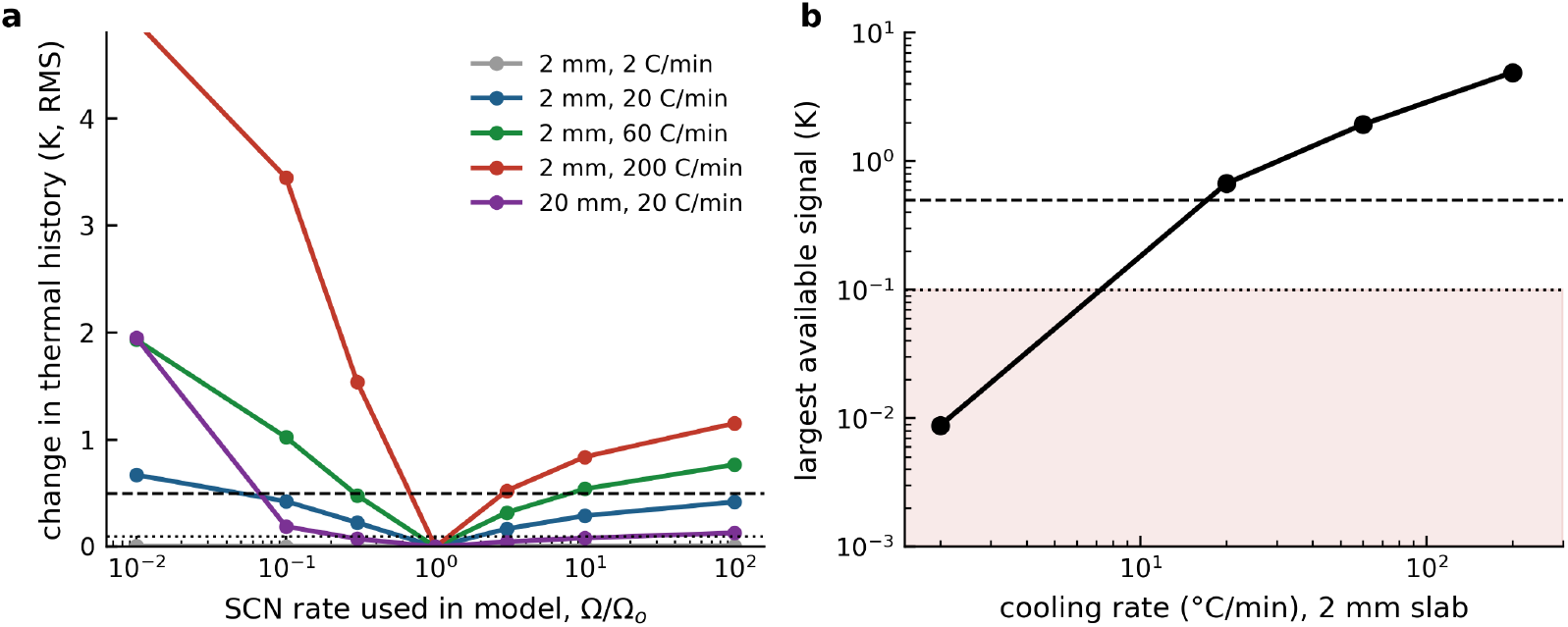
This figure demonstrates the cooling conditions under which a theoretical measurement methodology of thermal history during freezing of tissues can be used to extract by fitting the underlying unknown IIF nucleation parameters. **Fig. 7(a)** shows the change in the predicted thermal history, as a root-mean-square over five probes, when the surface nucleation (SCN) rate used in the model is moved away from its measured value, for five different cooling protocols. The two dotted horizontal lines mark an anticipated thermocouple noise of either 0.1 or 0.5 K in the measurements. An inverse measurement methodology can only recover what the tissue responds to as exhibited in the measured data. Our synthetic simulations suggest that at a cooling rate of 2 °C/min the entire range from no IIF whatsoever to a hundred-fold rate increase only manifests in the measured thermal history as 0.01 K change. This is because the cell is already 97 per cent dehydrated when the IIF processes become active and so has almost no latent heat left to release (and subsequently, does not at all impact the measured temperature values). However, raising the cooling rates to 20, 60 and 200 °C/min lifts the available signal to 0.67, 1.93 and 4.90 K, respectively and all of which are above the expected inherent thermocouple error of 0.5 K. This is an obvious result and shows that the IIF constants become “measurable” in the same regime in which they begin to “expend” a measurable amount of latent heat. **Fig. 7(b)** shows the largest change available signal change (in K) anywhere in the tested range for various cooling rates for the 2 mm slab. The shaded band lies below the resolution of a fine thermocouple or an anticipated error of 0.1 K error in measurements. The higher dotted line represents a more realistic thermocouple measurement error of 0.5 K and the smallest usable cooling rate of ~20 °C/min on a 2 mm slab to realistically be able measure the thermal history to inversely recover the IIF nucleation parameters via well-established curve-fitting procedures.

Figs. 8a and 8b show that treating a noisy coupled solution at nominal parameters as the measurement, and scanning both constants, returns the true values at the grid minimum, with a misfit equal to the noise floor and 1 to 5 K for wrong parameters. The recoverable region is narrow in Ω_*o*_ and elongated in *κ*_*o*_ or ±26% within one grid step and 0.93 to 1.06 of the true *κ*_*o*_ at a measurement noise of 0.1K. This widens to 0.59 to 1.41 within one grid step and 0.5 to 2.0 of true *κ*_*o*_ at a measurement noise of 0.5K. So, *κ*_*o*_ is now constrained only to a factor of 2. The inverse fitting also found no compensating ridge, i.e., the best-fit Ω_*o*_ does not move when *κ*_*o*_ is mis-specified. This is consistent with the two constants controlling different features of the SCN formation as *κ*_*o*_ controls the temperature at which the latent heat appears while Ω_*o*_ determines how abruptly the latent heat manifests in the formulation. Above roughly ten times the true Ω_*o*_ the misfit plateaus, so a sufficiently fast true rate would yield a lower bound rather than a true and accurate value. It is important to state that these results establish identifiability and feasibility of the inverse problem within the model and not the measurability or feasibility in a laboratory.

**Fig. 8.**
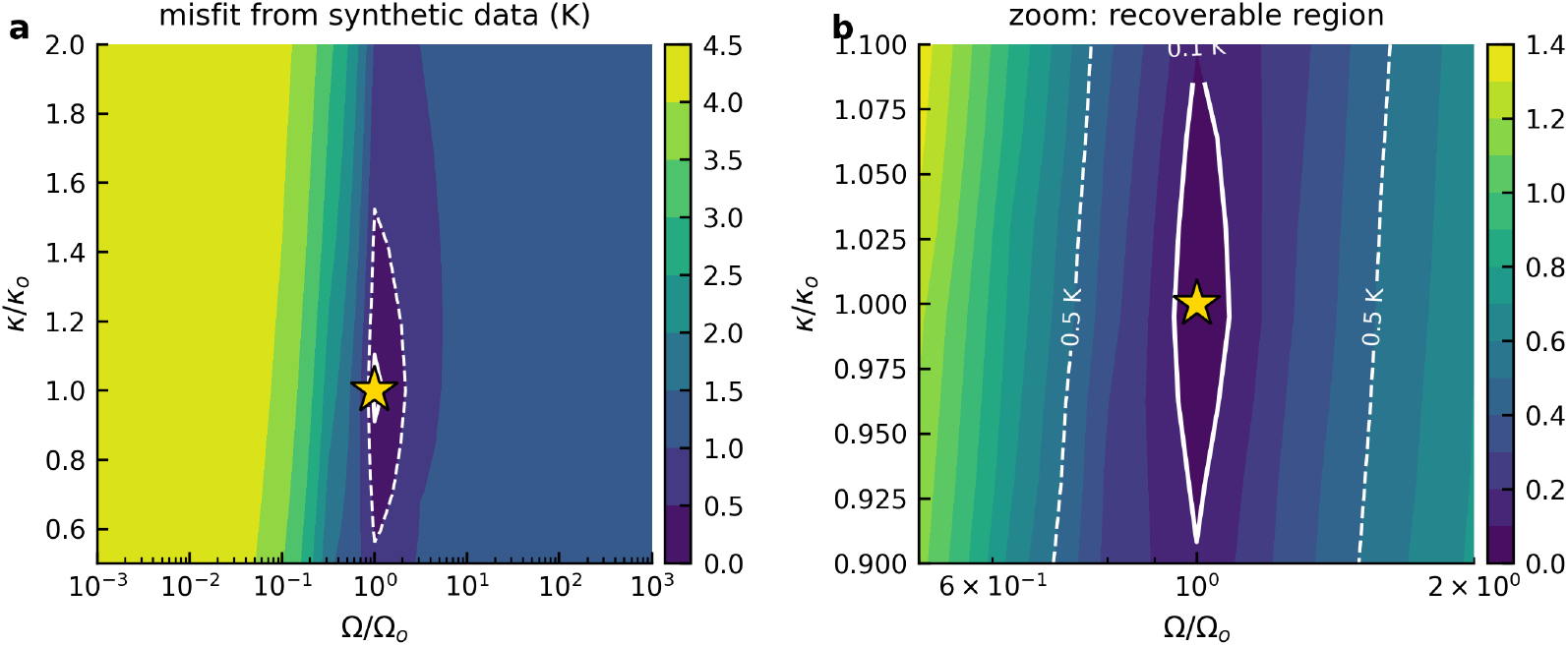
This figure demonstrates the errors associated with the inverse fitting procedure and the unequal degree of uncertainty associated with recovering the underlying SCN IIF parameters (Ω_*SCN*_ and *κ*_*SCN*_). Specifically, the “misfit” between trial parameters and a synthetic measurement, which is a coupled solution at the measured constants for the 2 mm slab at 200 °C/min with 0.1 K Gaussian noise added, is shown as a contour plot. The quantity plotted is the model separation or the part of the misfit not due to noise, see text for further details. Both graphs show the same data with **Fig. 8(b)** showing a “zoomed” in region near the “true” values of Ω_*SCN*_ and *κ*_*SCN*_ while **Fig. 8(a)** shows the “coarser” scan over six decades in the rate constant, Ω_*SCN*_ and a factor of four in the exponent constant, *κ*_*SCN*_ for 117 coupled runs. The white contours in the figures mark either the 0.1 or the 0.5 K error in the thermocouple measurements. The star represents the ground truth or the true value of Ω_*SCN*_ and *κ*_*SCN*_. The minimum “misfit” values lies at the true parameters with a value at the noise floor (either 0.1 or 0.5K), while wrong parameters give errors up to 5.0 K. Note that the contour valley is narrow in the rate constant, Ω_*SCN*_ and elongated in the exponent constant, *κ*_*SCN*_. This suggests an inherent variability in our projected confidence of the recovered Ω_*SCN*_ and *κ*_*SCN*_ parameters with Ω_*SCN*_ having a higher degree of confidence (due to the narrower range) and conversely, *κ*_*SCN*_ has a lower degree of confidence (due to the elongated nature of the contour). The specific details are provided in the text. Also, the lack of a noticeable compensating ridge is consistent with the two constants controlling different features of the ice nucleation process. Outside a window of about three decades in the SCN rate the separation again changes by less than the thermo-couple noise, so rates far above or below the measured value are recoverable only as bounds and not as exact values.

Figs. 9a shows that driving a single cell with each protocol’s temperature history shows that in a 2 mm slab cooled to −80 °C, SCN always fires first at −10 °C for a cooling rate of 20 °C/min and at −13 °C for a cooling rate of 200 °C/min. Conversely VCN requires roughly 35 K of supercooling and is reached only at −40 °C, as demonstrated earlier, or by which time the cell has already nucleated via SCN. Only the fastest cooling rates of liquid-nitrogen quench inverts this, with SCN never reaching 50% and VCN firing at −40 °C with 99% of the cell water still unfrozen (Fig. 9b). A controlled-rate protocol held above −30 °C is therefore a two-parameter problem in SCN alone, i.e., we are determining only two parameters (Ω_*SCN*_and *κ*_*SCN*_) rather than four (Ω_*SCN*_, *κ*_*SCN*_, Ω_*VCN*_ and *κ*_*VCN*_). These methods suggest that, in theory, the parameters governing SCN IIF and VCN IIF in tissues can be extracted from measured temperature-time thermal history under appropriately chosen cooling conditions. However, experimental validation is necessary to lend credence to this assertion, as the simulations only demonstrate computational identifiability and motivate an experimental inverse-measurement strategy.

**Fig. 9.**
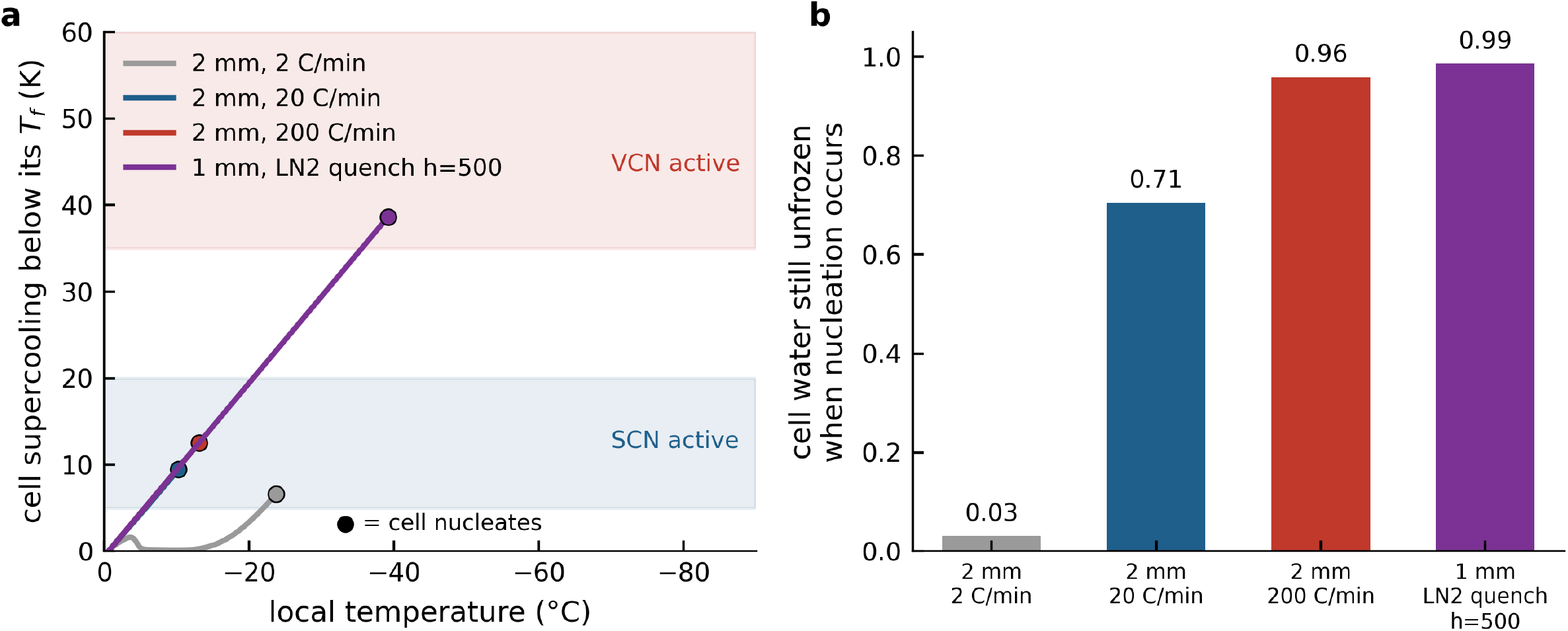
This figure demonstrates the possible scenarios under which the SCN and VCN IIF parameters can be recovered from the synthetic measurements. **Fig. 9(a)** shows the supercooling below its own freezing point experienced by a single cell driven by each protocol’s temperature history, plotted only while the cell is still liquid and able to exhibit intrcellular ice formation (see text for further details). The filled circles mark the nucleation. The lower shaded band is the range over which the SCN pathway is active and the upper band represents the VCN pathway active zone. Note that, in theory, the cooling protocols to “trigger” SCN and VCN are distinct enough that it should be possible to extract one set of parameters (either Ω_*SCN*_ and *κ*_*SCN*_ or Ω_*VCN*_ and *κ*_*VCN*_) without knowledge of the other, a less intractable problem than fitting for all 4 parameters from one data set. **Fig. 9(b)** shows the fraction of the cell’s water still unfrozen at the moment it nucleates, with the value printed above each bar. Until a cell dehydrates its supercooling tracks the local temperature, so the three faster protocols follow a common line, and only at a slow cooling rate of 2 °C/minute does dehydration keep pace and hold the supercooling near 6 K. In a 2 mm slab cooled to −80 °C the SCN pathway always fires first, at −10 °C for a cooling rate of 20 ° C/min and at −13 °C for a cooling rate of 200 °C/minute. Correspondingly, the VCN pathway needs about 35 K of supercooling and is reached only near −40 °C, after the cell has already frozen. Holding such a protocol above −30 °C therefore isolates the SCN pathway and reduces the inverse problem to two unknowns (Ω_*SCN*_ and *κ*_*SCN*_). The liquid nitrogen quench is the mirror image, with the SCN pathway never reaching half probability and the VCN pathway firing at −40 °C with 99% of the cell water still unfrozen. The trade-off inside the usable window is also shown as more retained water means a “stronger” measurement signal but at the cost of moving towards the VCN phenomena and away from the SCN one.

## DISCUSSION

The central quantity identified here, ∏ = *τ*_*osm*_/*τ*_*rec*_, compares the rate at which a cell can lose water through osmotic transport with the rate at which the local temperature traverses the phase change window. This competition is not a new concept and is inherent in the physical basis of Mazur’s two factor hypothesis [4], which requires cooling sufficiently slowly for cells to remain in chemical potential equilibrium with the extracellular solution and thereby avoid intracellular ice formation. The classical inverted U survival curve and the observation that optimal cooling rates can vary by several orders of magnitude among cell types, in accordance with differences in membrane permeability and surface to volume ratio, both follow from this underlying timescale competition. The present study extends this concept by establishing a quantitative relationship between the latent heat released during tissue freezing through microscale processes and the heat release predicted by the macroscale equilibrium phase diagram. These two processes are intrinsically coupled, leading to an important structural connection between cellular injury and model validity. Cells are most susceptible to intracellular ice formation under cooling conditions where the equilibrium enthalpy model is also most likely to become inaccurate. In other words, a protocol operating safely within Mazur’s slow cooling regime is also a regime in which the enthalpy formulation remains reliable. As cooling conditions shift toward the rapid cooling regime, both protections can be lost simultaneously. Two independent observations support this interpretation. The first is the equilibrium identity shown in Fig. 2(a), which explains why the coupled and uncoupled formulations converge in the quasi static limit. The second is the extremely large value of *τ*_*SCN*_, approximately 10^1−2^ s, at 2 K of supercooling. This provides a physical basis for Mazur’s empirical observation that cells experiencing approximately 2 K of supercooling are unlikely to nucleate intracellularly [4,7]. The fixed width of the lag band identified in this study, shown in Fig. 2(b), also indicates that discrepancies between coupled and uncoupled models should always be interpreted in relation to the relevant time and length scales. The lag width scales with the product of the freezing front velocity and *τ*_*osm*_, whereas the frozen depth increases with the product of the freezing front velocity and time. Consequently, the relative error decreases as freezing progresses and the characteristic frozen depth becomes larger. Larger domains can therefore conceal the underlying discrepancy even when the absolute lag width remains essentially unchanged. This scale dependence may help explain why an effect of this magnitude has received relatively little attention in previous tissue freezing studies.

Stated in the language of computational heat transfer rather than cryobiology, the significance of the ∏ grouping lies in determining the validity of an equilibrium closure for a source term. Enthalpy and apparent heat capacity methods are widely used for phase change problems [47,57–86,101–110]. These approaches assume that latent heat is released locally as the temperature crosses the phase change interval, which implicitly assumes that the microscale processes responsible for releasing that heat are effectively instantaneous relative to the evolution of the thermal field. This assumption is often made implicitly and is rarely evaluated quantitatively. The dimensionless group derived here can therefore be interpreted as a local Damköhler number for this equilibrium closure. It compares the relaxation time of the subgrid process with the residence time of the thermal field within the temperature range over which the source term is active. When ∏ is small, the equilibrium closure is valid and the uncoupled enthalpy formulation provides an accurate representation of the coupled system. As ∏ approaches unity, the source term increasingly lags behind the thermal field that drives it, producing a growing discrepancy between the coupled and uncoupled formulations. The error eventually approaches a saturation value determined by the amount of enthalpy that the subgrid process can temporarily withhold. The saturation limit predicted from tissue properties alone and confirmed by the simulations provides a quantitative manifestation of this behavior. Viewed in this broader framework, the finding is not specific to biological tissue. The same structure can arise in any phase change problem in which a finite-rate microscale process controls the timing of latent heat release. Alloy solidification provides a close analogue [111–117]. In that setting, equilibrium partitioning and the lever rule represent one limiting case, while the opposite limit of negligible back-diffusion represents another. The actual behavior depends on the competition between transport and cooling timescales. Similar microscale transport limitations arise in food freezing, phase change thermal energy storage, and freezing in porous media [118–125], where continuum formulations often treat microscale processes as instantaneous. The cryobiological setting offers a particularly useful opportunity because the relevant subgrid processes are sufficiently well characterized that their timescale competition can be calculated rather than inferred. This makes it possible to derive a quantitative criterion for determining when an equilibrium closure is valid and when finite-rate microscale physics must be explicitly resolved.

The definition of *τ*_*osm*_ reveals three distinct physical scaling relationships, each of which is consistent with physical intuition. First, 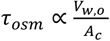, which scales directly with the characteristic cell length. Thus, a cell that is twice as large in every dimension requires approximately twice as long to reach osmotic equilibrium. Second, 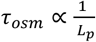, which reflects the expected behavior of membrane permeability. A more permeable membrane allows water to move more rapidly and therefore reduces the osmotic relaxation time. Because *L*_*p*_ decreases steeply with decreasing temperature according to the Arrhenius relationship defined above, *τ*_*osm*_ increases as the cell cools and the membrane becomes progressively less permeable. Third, 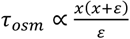, which is less intuitive but has an important physical interpretation. This factor increases sharply as *x* approaches one, corresponding to a cell that retains most of its intracellular water. Consequently, a cell with a small *x*, corresponding to substantial dehydration, relaxes much more rapidly than a cell that remains highly hydrated. In other words, osmotic relaxation becomes progressively easier once the cell has begun to dehydrate, while the slowest response occurs at the beginning of the process when the cell is still close to fully hydrated. This behavior explains the spatial distribution of the discrepancy between the coupled and uncoupled models. The largest disagreement is concentrated near the advancing freezing front, where cells remain close to their initial hydrated state. Farther behind the freezing front, cells have had more time to dehydrate and shrink, which reduces the osmotic relaxation time and allows the coupled and uncoupled solutions to converge.

The comparison presented in Fig. 3 is, to our knowledge, the first direct evaluation of the timescale against which *τ*_*osm*_ competes in a tissue-scale freezing calculation. The result is clear. A dimensionless group based on domain-scale thermal diffusion provides only weak organization of the data, whereas a group based on local residence time consistently orders the results across a broad range of conditions. This distinction has direct implications for extrapolating tissue freezing results across scales. Cryobiological scale-up is conventionally framed in terms of characteristic diffusion lengths, and the intuition that larger systems are more challenging is well established for thermal uniformity, cracking, and cryoprotectant delivery. For the validity of the latent-heat treatment, however, the dependence is reversed. At a fixed cooling rate, larger domains produce longer local residence times and therefore smaller values of ∏. The failures identified here occur primarily in small samples and in samples cooled simultaneously from multiple directions. For example, between the operating point examined by Devireddy et al. [1] and the failure threshold at ∏ = 0.1, *τ*_*osm*_ remains unchanged at approximately 24 s because it is a tissue property, whereas *τ*_*res*_decreases from approximately 495 s to less than 40 s. No unusual cellular properties are therefore required to reach the failure regime. This explains why an 8 mm cube cooled at 20 °C/min, which represents an entirely routine protocol, can fall on the coupled side of the model validity boundary.

That intracellular ice formation releases latent heat is well established. However, to the best of our knowledge, the observation that IIF can restore the accuracy of a model that neglects cellular kinetics and microscale phenomena has not previously been recognized. This finding has an important practical implication that runs counter to standard cryopreservation practice. Suppressing intracellular ice formation is a primary objective of cryopreservation protocol design because IIF is generally considered harmful and potentially lethal to cells [4,7,8,10,21,27,28,30,45–47,71,78,82–86]. A protocol that successfully suppresses IIF through increased cryoprotectant loading, enhanced membrane permeability, or operation under conditions where nucleation kinetics are slow moves the system toward the upper curve in Fig. 6(a), where the enthalpy model used to design the protocol becomes least reliable. Conversely, this framework may help explain why enthalpy-based predictions have been effective in cryosurgery, where protocols deliberately drive tissue toward complete intracellular freezing and therefore maintain the latent heat contribution from IIF. The presence of VCN in addition to SCN introduces two distinct nucleation pathways and unexpectedly makes the single-parameter lag law more robust. Including both pathways improved the fit across the full dataset, reducing the residual from 17.5 K to 8.1 K. It also relaxes the design criterion shown in Fig. 6(b) rather than tightening it. For example, at a 3 K error tolerance, ∏ _*to*I_ increases from 0.102 for SCN alone to 0.148 when both SCN and VCN are included. Thus, incorporating the more complete physics allows the enthalpy model to remain valid over a wider range of conditions than the model containing only SCN. This behavior is opposite to the usual expectation that adding a previously neglected mechanism will narrow the validity range of a reduced model.

The most consequential current application of tissue-scale heat transfer modeling is organ banking by vitrification, where cooling and rewarming must exceed critical rates while remaining sufficiently uniform to avoid fracture [50,51,54–56,68]. Recent demonstrations include recovery and transplantation of rat kidneys following vitrified storage with restored function [13], vitrification and nanowarming of rat livers [12], and physical vitrification of multi-liter volumes and whole porcine liver tissue [126,127]. Critical cooling rates for current cryoprotectant formulations are on the order of a few °C/min, while critical warming rates are typically in the tens to hundreds of °C/min. Typical convective boundary conditions correspond to film coefficients on the order of 100 W m^−2^ K^-1^, while controlled-rate freezers can achieve cooling rates of tens of °C/min. These conditions fall within the parameter ranges explored in the present study. We emphasize that this connection should not be overstated. Successful vitrification is, by definition, an ice-free process, whereas the ∏ formalism developed here describes a freezing system. Importantly, the present model does not include cryoprotectants. The relevance to vitrification therefore lies in the broader modeling methodology rather than in vitrification itself. At organ scale, protocol design relies on continuum heat transfer simulations to determine whether local cooling and warming rates exceed critical values throughout heterogeneous tissue. If vitrification fails or partial crystallization occurs, however, latent heat released during ice formation can feed back into the thermal field and potentially introduce some of the same multiscale limitations identified in the present study. The precise nature of this regime and its implications for cryoprotectant-loaded systems remain topics for future investigation.

Our model does not account for the presence of cryoprotective agents (CPAs), which can alter cell membrane transport properties, at least in isolated cells in suspension [6,29,46,47,71,128–139]. For example, in isolated rat hepatocytes, exposure to 0 to 2 M DMSO reduces *L*_*pg*_ by approximately a factor of six, while *E*_*Lp*_ is reduced by approximately a factor of three [29]. Because *τ*_*osm*_is inversely proportional to *L*_*p*_, a cursory analysis suggests that, under otherwise similar cooling conditions, the presence of DMSO would increase ∏ and could shift the system into the regime where the coupled model is required. A related but distinct limitation concerns the tissue-level water transport parameters themselves. The calorimetric technique used to measure *L*_*pg*_ and *E*_*Lp*_ is constrained by instrument capabilities and is currently applicable only to cooling rates at or below approximately 40 °C/min [35,36]. More broadly, ice nucleation parameters measured in isolated cells may not necessarily translate to intact tissue. Preliminary unpublished measurements from our group, obtained using an extension of the previously described calorimetric method, suggest that ice nucleation kinetics in whole rat liver tissue differ by more than 30% from those measured in isolated hepatocytes. Incorporating tissue-level measurements into the present framework would shift the ∏ axis in Figs. 3, 4, and 6, but would not change the overall qualitative conclusions.

Several assumptions from the original model are retained in the present formulation. These include the Krogh cylinder representation, constant membrane surface area, and instantaneous equilibration of the extracellular phase. The assumption of instantaneous extracellular equilibration could be relaxed in future models to determine whether extracellular transport becomes rate limiting in dense tissue matrices with complex and convoluted vasculature. The present formulation also does not account for tissue mechanics. Thermal and osmotic loading during freezing can generate mechanical stresses, and freezing-induced cracking has been documented at the organ scale [54–56,68]. At the cellular scale, growing extracellular ice crystals may also interact mechanically with cells trapped within unfrozen channels [140,141]. These effects are beyond the scope of the present thermodynamic formulation. In particular, the latter effects relate to the unfrozen fraction mechanism proposed by Nei and incorporated into Mazur’s framework. Recent evidence suggests that ice crystal spacing alone can cause cellular disruption independently of the osmotic and thermal effects considered here [140,141]. The present analysis is also restricted to cooling and does not consider rewarming processes such as recrystallization and devitrification [52]. Finally, ∏ is evaluated at a single reference state, while the residence time is represented by the median value across probe locations. The resulting criterion therefore characterizes typical rather than worst-case behavior. For conservative model selection, the shortest relevant local residence time should instead be used. Nevertheless, the range of approximately ∏ = 0.2 to 3, which encompasses the routine protocols shown in Fig. 5, remains the most strongly supported regime for the present criterion.

Another limitation concerns the temperature range from −5 to −20 °C over which SCN parameters are typically determined [18–25,28,76–86], as well as the use of saline droplets as biological surrogates for estimating VCN parameters [87]. These measurements may not accurately represent the corresponding parameters in intact tissue, and the associated uncertainty is greatest under liquid nitrogen quench conditions where ∏ > 40. Accordingly, the results presented for ∏ > 40 should be interpreted as bounds rather than as a quantitative representation of the true tissue response. The SCN-only simulations and the combined SCN-VCN simulations shown in Fig. 4 bracket this uncertainty. Given the uncertainty surrounding IIF parameters and the limited availability of methods for measuring them directly, we explored and validated an inverse problem approach, as described in the Results. Our analysis indicates, however, that careful consideration must be given to the cooling conditions under which this approach can be applied. In particular, the inverse problem is most informative in the regime where microscale transport significantly influences the macroscale tissue freezing response. Consequently, the experiment must be performed under conditions where the underlying physics is most complex and the associated computational cost is highest. The sensitivity of the proposed measurement also saturates at both extremes of the parameter range. At approximately ten times the baseline value of Ω_*SCN*_, the predicted thermal history becomes largely insensitive to further increases in nucleation rate, changing by only approximately 0.2 K per decade of nucleation rate. This change is comparable to thermocouple noise. Thus, when the true nucleation rate is sufficiently high, the inverse approach can recover only a lower bound on the parameter. Conversely, below approximately one hundredth of the baseline value, the thermal history again becomes insensitive to the nucleation rate because nucleation no longer contributes significantly to latent heat release. Under these conditions, the true nucleation parameter can be recovered only as an upper bound. Between these two limits, over an intermediate range spanning approximately three decades, the parameter recovery was unique. The misfit exhibited a well-defined minimum at the true parameter values, with no compensating ridge between Ω_*SCN*_and *κ*_*SCN*_. These results define the practical operating window for the proposed inverse measurement approach and highlight the importance of selecting experimental conditions within the regime where the thermal history remains sufficiently sensitive to the underlying intracellular ice formation parameters.

The SCN/VCN IIF parameters identified with our inverse problem strategy will be sensitive to any errors in and the choice of all the other model parameters (*L*_*pg*_, *E*_*Lp*_, Krogh cylinder dimensions, *f*_*ic*_, the thermophysical properties, boundary conditions, and so on). Given the sensitivity of ice nucleation to the specified heat transfer coefficient, our recommendation is to perform the experiments at a constant temperature (Dirichlet) boundary conditions and to propagate the published uncertainties in *L*_*pg*_ and *E*_*Lp*_ through the inversion rather than treating them as exact. An important limitation of the inverse analysis is that the synthetic measurements were generated from the same model used for parameter recovery. The exercise therefore demonstrates computational identifiability rather than experimental validation. Measurement noise was introduced to approximate experimental uncertainty, but independent experimental data will be required to establish whether the proposed approach can recover tissue-level nucleation parameters. Future studies will explore this limitations both computational and experimental to test whether protocols of different severity return consistent constants, and to the possibility of extending the design to the deep-quench regime where VCN parameters can be isolated in the same way.

Future studies should extend the present analysis to cryoprotectant-loaded systems, which is an important next step toward organ-scale relevance. This extension is straightforward within the current framework and requires incorporation of a ternary phase diagram together with concentration-dependent transport parameters. The same approach is already being applied to the design of oocyte cryopreservation protocols [23,26,74,142]. Recalibrating the VCN parameters directly for biological cells would further constrain the quench-regime boundary rather than simply broadening it, and such recalibration is within reach of existing experimental methods [28,33]. The proposed criterion is also experimentally testable without specialized equipment. It predicts differences of tens of percentage points in frozen volume between millimeter-scale samples cooled from a single face and those cooled from multiple faces at identical nominal cooling rates. This comparison could be performed using a controlled-rate freezer together with imaging of the advancing frozen boundary. More broadly, increasingly detailed microscale descriptions of coupled water transport and heat transfer in cell and tissue cryopreservation are becoming available, along with agent-based models of intracellular ice nucleation and propagation through tissue [143–150]. These models are considerably more computationally expensive than continuum formulations. This trade-off is not unique to cryobiology and also arises in other systems where thermal gradients are coupled to stochastic phase change processes, such as the freezing step in pharmaceutical freeze-drying [151–156]. A criterion analogous to the ∏ grouping identified in the present study could therefore provide a valuable design tool for determining in advance when additional microscale physics will alter the macroscale response and when it will not. Such a criterion would enable computationally intensive models to be deployed selectively under conditions where they are needed, rather than applied uniformly, thereby improving both computational efficiency and model selection for multiscale freezing problems.

## CONCLUSION

We revisit the assertion of Devireddy et al. [1] that microscale freezing processes do not alter the macroscale tissue freezing process and show that, although this assertion is valid under the conditions examined in the original analysis, it does not hold universally. Using an appropriate scaling analysis, we identified a non-dimensional grouping, ∏ = *τ*_*osm*_/*τ*_*rec*_, that organizes a broad range of simulation data and provides a criterion of ∏ > 0.1for determining when the coupled biophysical model of tissue freezing is required. Five key findings emerge from this analysis. First, the discrepancy between the coupled and uncoupled formulations is purely kinetic, with the associated lag zone occupying a band of approximately fixed width at the advancing ice front. This behavior can mask the true magnitude of the model error because the relative error decreases with time as the ice front advances. Consequently, discrepancies reported across different studies cannot be meaningfully compared without specifying the relevant time and length scales. Second, substantial divergence between the two models requires both suppressed ice nucleation and ∏ > 0.1. These conditions imply that cooling protocols designed to suppress intracellular ice formation are also those most likely to require consideration of the underlying microscale biophysical processes. Neglecting these processes under such conditions can therefore lead to erroneous predictions. Third, cooling geometry alone can shift a protocol between the uncoupled and coupled regimes. For example, cooling an 8 mm cube from a single face can be adequately described using the uncoupled formulation, whereas cooling the same cube simultaneously from all six faces necessitates the use of the coupled micro-macroscale model. Fourth, the closed-form criterion provides a user-calibrated design tool for selecting the appropriate tissue freezing model under a prescribed set of cooling conditions and an acceptable error tolerance. Fifth, our inverse analysis indicates the possibility of extracting intracellular ice formation parameters from macroscale thermal history measurements within a narrow range of experimental conditions. If experimentally validated, this approach could enable bioengineers to measure the SCN and VCN intracellular ice formation parameters in intact tissue for the first time. Such a capability could represent a significant advance in cryobiological measurement and characterization.

## Appendix A. The 124 one-dimensional cases

Blank input cells mean the parameter was held at its rat liver reference value: L_pg = 3.0e-13 m3 N-1 s-1, E_Lp = 265 kJ/mol, f_ic = 0.589, cell size 1x (dx 22 um, r_vo 3.8 um, l 11.4 um), Omega_SCN = 8.5e9 m-2 s-1. dT_max is the largest thermal-history difference between the coupled and uncoupled models over all probes and times. The front-arrival error is negative because the coupled front always arrives first. All values use the converged Picard settings (tolerance 1e-8, up to 200 iterations).

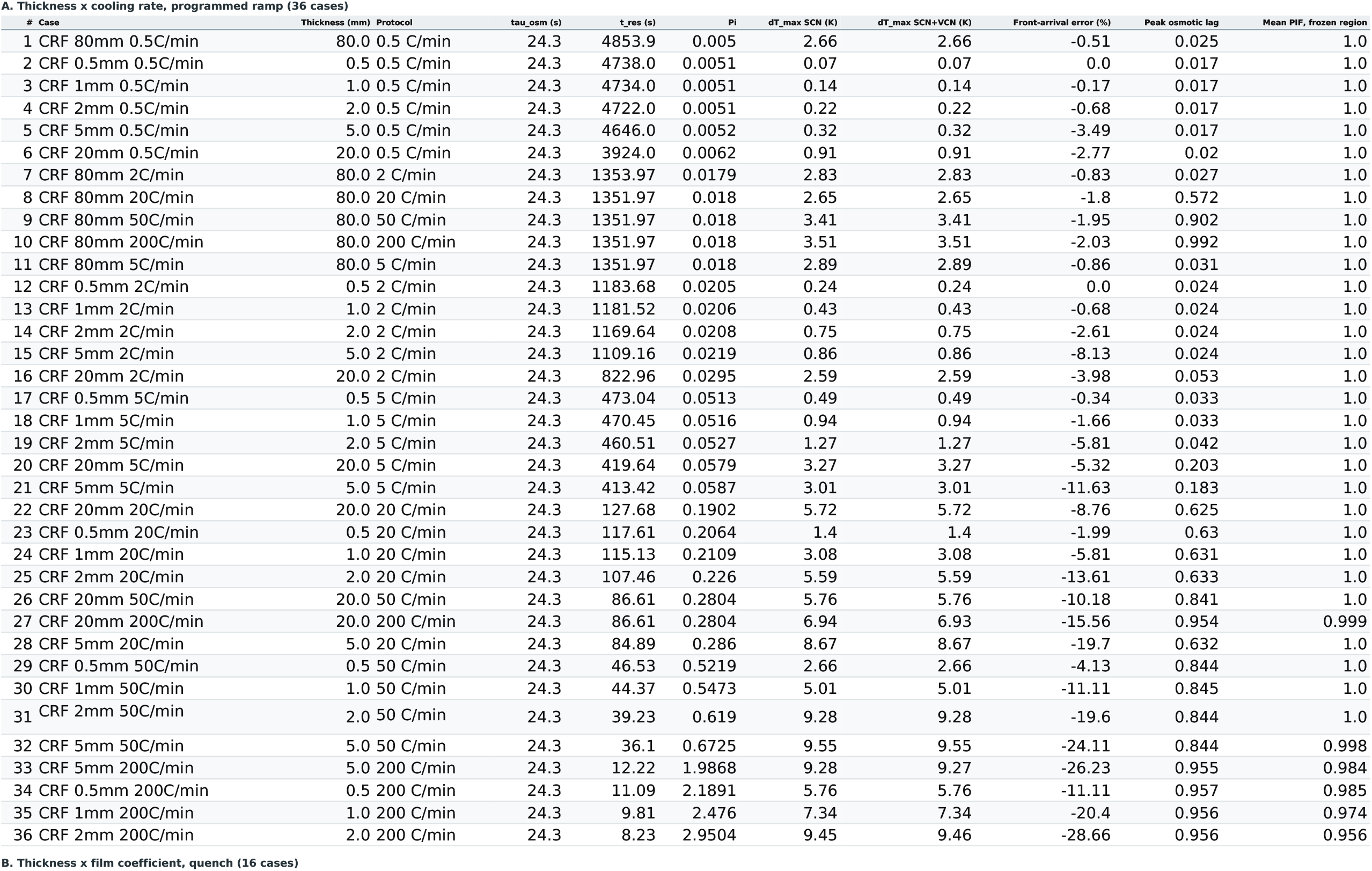
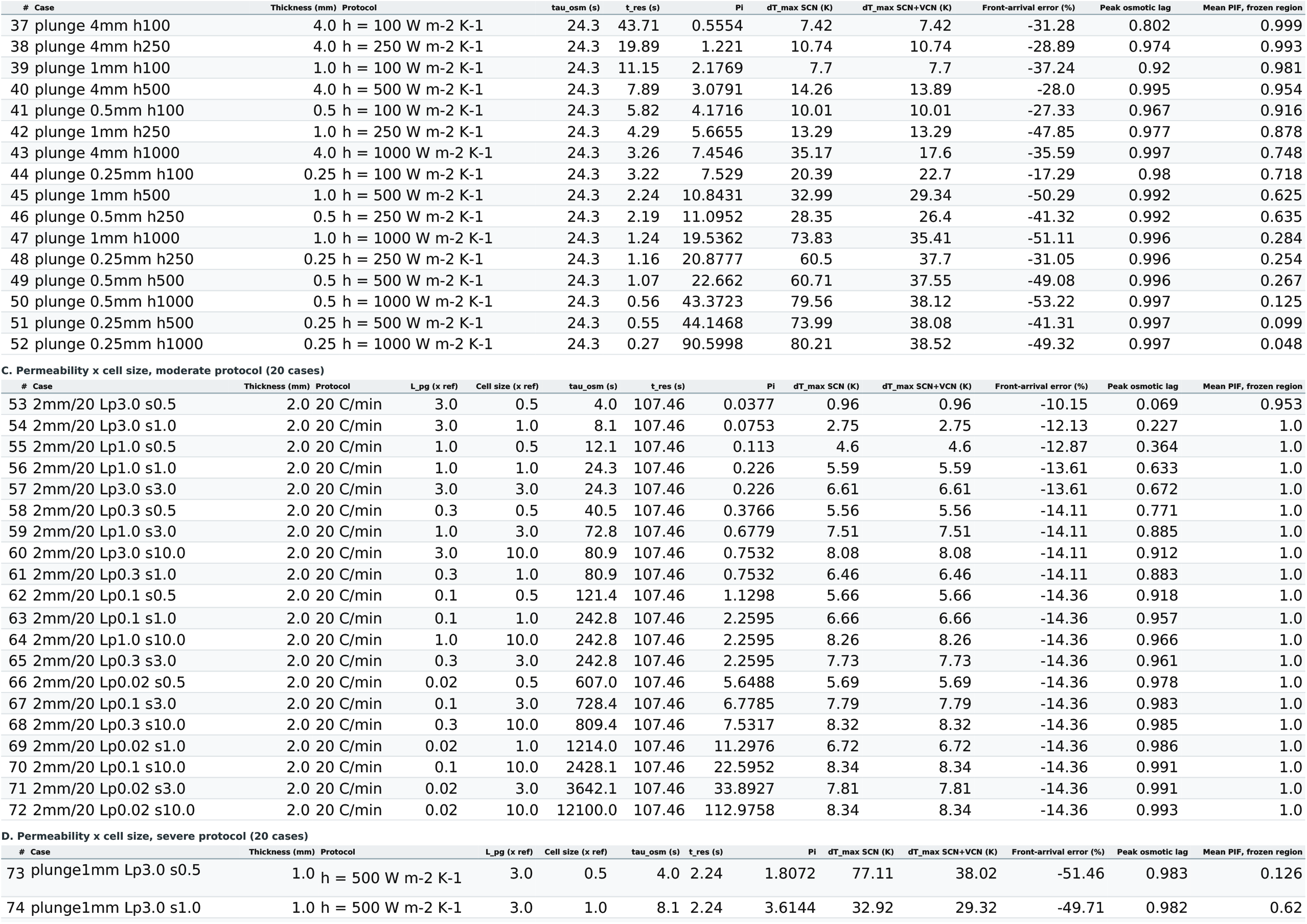
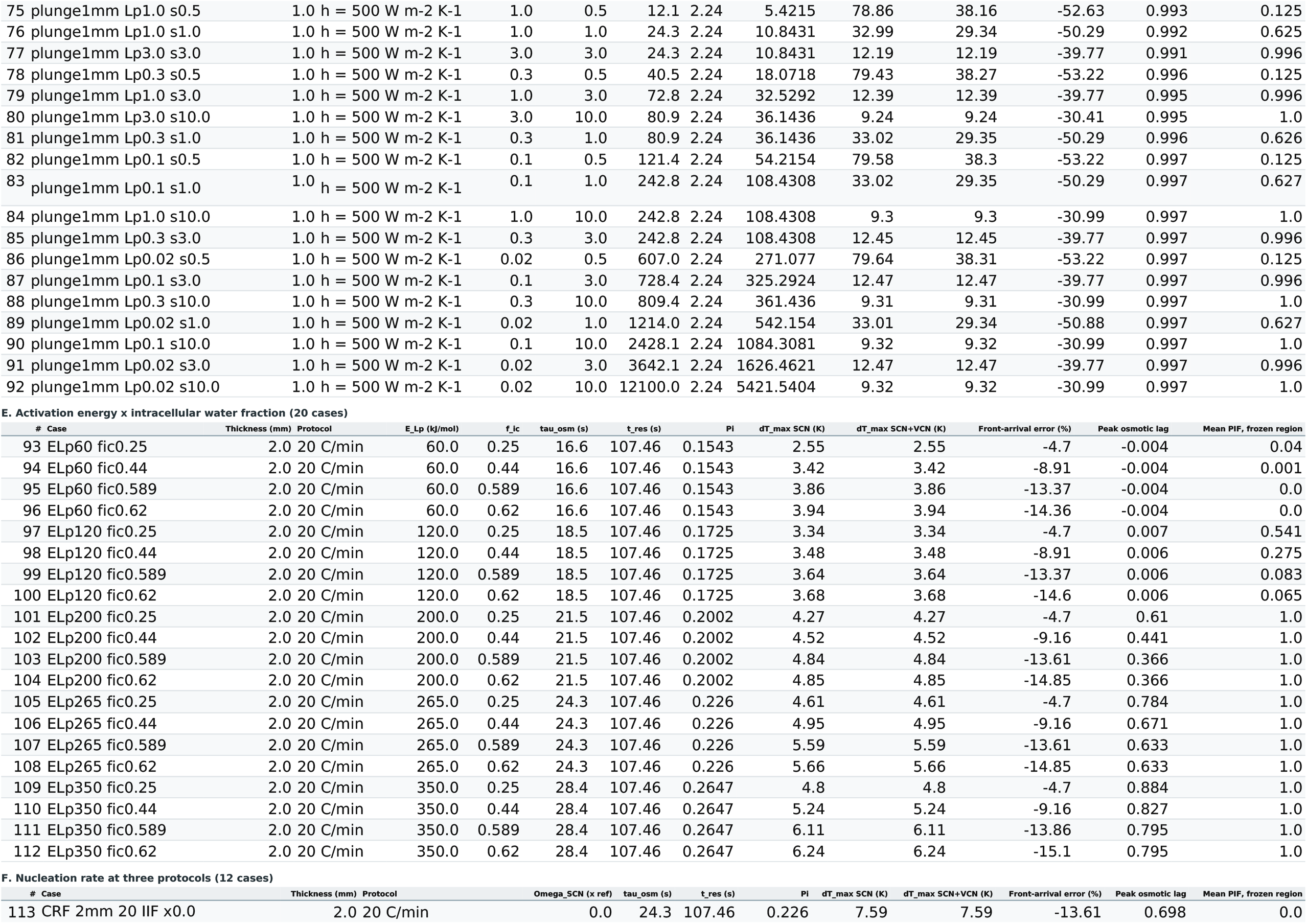
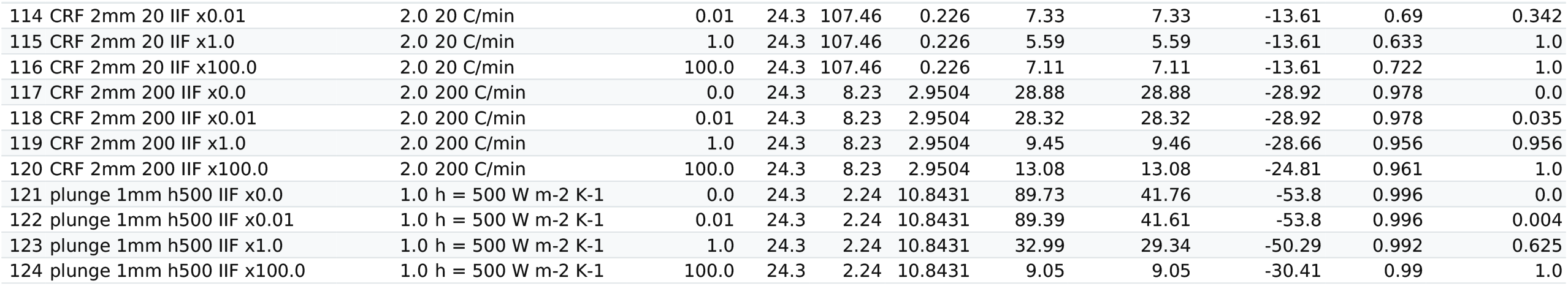

## Appendix B. The 42 nucleation-window cases

These cases hold the protocol fixed and vary only the surface nucleation rate, so each shares its coupling group with the parent case in Appendix A. They are not binned in Table 3 for that reason. They support Fig. 4(b) and the suppressed-nucleation curve of Fig. 6.

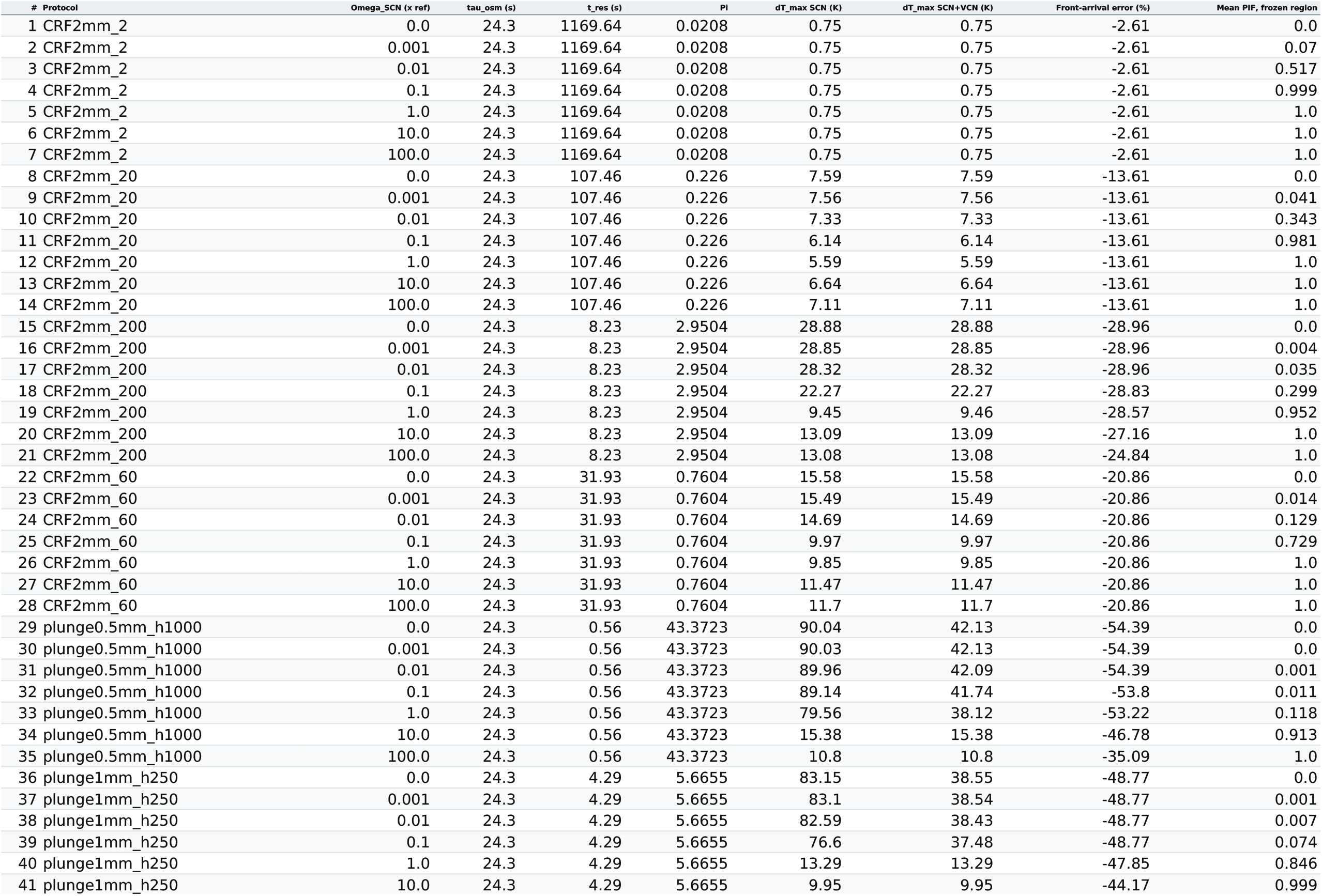
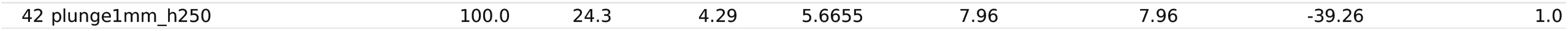

## Appendix C. The 6 two- and three-dimensional cases

Frozen-volume gap is the largest difference between the two models in the fraction of the domain below the phase-change temperature, in percentage points. Thermal errors carry roughly 12 per cent numerical uncertainty from solver settings; the frozen-volume gaps changed by at most 0.11 points under the same test.

| # | Case | Grid | tau_osm (s) | t_res (s) | Pi | dT_max SCN (K) | dT_max SCN+VCN (K) | Frozen-volume gap SCN (pts) | Frozen-volume gap SCN+VCN (pts) |
| --- | --- | --- | --- | --- | --- | --- | --- | --- | --- |
| 1 | 2D-A_cryoprobe | 110x110 | 24.3 | 73.5 | 0.33 | 4.67 | 4.67 | 1.8 | 1.8 |
| 2 | 2D-B_block_corner | 100x100 | 24.3 | 36.0 | 0.674 | 7.41 | 7.41 | 38.3 | 38.3 |
| 3 | 2D-C_vitrification | 80x80 | 24.3 | 0.25 | 97.123 | 71.29 | 34.55 | 39.6 | 39.6 |
| 4 | 3D-A_cube8mm_20Cmin | 40x40x40 | 24.3 | 110.0 | 0.221 | 6.96 | 6.96 | 25.4 | 25.4 |
| 5 | 3D-B_cube4mm_60Cmin | 40x40x40 | 24.3 | 36.75 | 0.661 | 5.56 | 5.56 | 32.8 | 32.8 |
| 6 | 3D-C_cube1mm_plunge | 36x36x36 | 24.3 | 0.18 | 131.247 | 70.76 | 44.22 | 31.7 | 31.7 |

## Footnotes

1 Every symbol that appears anywhere (and in the model) is listed here with its meaning and units. SI units are used throughout (meters, kilograms, seconds, Kelvin, joules) unless otherwise noted.

## REFERENCES

1. Devireddy RV, Smith DJ, Bischof JC. Effect of microscale mass transport and phase change on numerical prediction of freezing in biological tissues. J. Heat Transfer. 2002 Apr 1;124(2):365–74.

2. Polge C, Smith AU, Parkes AS. Revival of spermatozoa after vitrification and dehydration at low temperatures. Nature. 1949 Oct 15;164(4172):666.

3. Lovelock JE. The haemolysis of human red blood-cells by freezing and thawing. Biochim Biophys Acta. 1953 Mar;10(3):414–26.

4. Mazur P. Cryobiology: the freezing of biological systems. Science. 1970 May 22;168(3934):939–49. doi: 10.1126/science.168.3934.939.

5. Love RM. Structure of animal tissue after freezing. Curr Probl Clin Biochem. 1971;3:18–34.

6. Meryman HT. Cryoprotective agents. Cryobiology. 1971 Apr;8(2):173–83.

7. Mazur P. Freezing of living cells: mechanisms and implications. Am J Physiol. 1984 Sep;247(3 Pt 1):C125–42.

8. Diller KR. Engineering-based contributions in cryobiology. Cryobiology. 1997 Jun;34(4):304–14. doi: 10.1006/cryo.1997.2012.

9. Devireddy RV. Biopreservation: Heat/mass transfer challenges and biochemical/genetic adaptations in biological systems. Heat Transf Res. 2013;44(3-4):245–272.

10. Shaik SM, Devireddy R. Heat and mass transfer models and measurements for Low-Temperature storage of biological systems. Handbook of Thermal Science and Engineering. 2018:2417

11. Taylor MJ, Weegman BP, Baicu SC, Giwa SE. New Approaches to Cryopreservation of Cells, Tissues, and Organs. Transfus Med Hemother. 2019 Jun;46(3):197–215.

12. Sharma A, Lee CY, Namsrai BE, Han Z, Tobolt D, Rao JS, Gao Z, Etheridge ML, Garwood M, Clemens MG, Bischof JC, Finger EB. Cryopreservation of Whole Rat Livers by Vitrification and Nanowarming. Ann Biomed Eng. 2023 Mar;51(3):566–577.

13. Han Z, Rao JS, Gangwar L, Namsrai BE, Pasek-Allen JL, Etheridge ML, Wolf SM, Pruett TL, Bischof JC, Finger EB. Vitrification and nanowarming enable long-term organ cryopreservation and life-sustaining kidney transplantation in a rat model. Nat Commun. 2023 Jun 9;14(1):3407.

14. Khaydukova IV, Ivannikova VM, Zhidkov DA, Belikov NV, Peshkova MA, Timashev PS, Tsiganov DI, Pushkarev AV. Current State and Challenges of Tissue and Organ Cryopreservation in Biobanking. Int J Mol Sci. 2024 Oct 16;25(20):11124.

15. Kim S, Rahaman KA, Lee JH, Han HS, Jeon H, Kim Y. Materials strategies and multidisciplinary approaches for low-temperature preservation of 3D cellular systems. Biomed Eng Lett. 2025 Dec 29;16(2):237–250.

16. Kang XY, Cheng JY, Ge WY, Tong YM, Yin DC. Revolution in Organ Preservation: Technological Exploration. Acta Biomater. 2025 Jun 1;199:50–73.

17. Thirumala S, Devireddy RV. A simplified procedure to determine the optimal rate of freezing biological systems. J Biomech Eng. 2005;127(2):295–300.

18. Scheiwe MW, Körber C. Thermally defined cryomicroscopy and thermodynamic analysis in lymphocyte freezing. Cryobiology. 1984 Feb;21(1):93–105.

19. Schwartz GJ, Diller KR. Intracellular freezing of human granulocytes. Cryobiology. 1984 Dec;21(6):654–60.

20. Mazur P, Rall WF, Leibo SP. Kinetics of water loss and the likelihood of intracellular freezing in mouse ova. Influence of the method of calculating the temperature dependence of water permeability. Cell Biophys. 1984 Sep;6(3):197–213.

21. Scheiwe MW, Körber C. Quantitative cryomicroscopic analysis of intracellular freezing of granulocytes without cryoadditive. Cryobiology. 1987 Oct;24(5):473–83.

22. Shabana M, McGrath JJ. Cryomicroscope investigation and thermodynamic modeling of the freezing of unfertilized hamster ova. Cryobiology. 1988 Aug;25(4):338–54.

23. Toner M, Cravalho EG, Karel M, Armant DR. Cryomicroscopic analysis of intracellular ice formation during freezing of mouse oocytes without cryoadditives. Cryobiology. 1991 Feb;28(1):55–71.

24. Toner M, Cravalho EG, Stachecki J, Fitzgerald T, Tompkins RG, Yarmush ML, Armant DR. Nonequilibrium freezing of one-cell mouse embryos. Membrane integrity and developmental potential. Biophys J. 1993 Jun;64(6):1908–21.

25. Karlsson JO, Cravalho EG, Borel Rinkes IH, Tompkins RG, Yarmush ML, Toner M. Nucleation and growth of ice crystals inside cultured hepatocytes during freezing in the presence of dimethyl sulfoxide. Biophys J. 1993 Dec;65(6):2524–36.

26. Mazur P, Seki S, Pinn IL, Kleinhans FW, Edashige K. Extra- and intracellular ice formation in mouse oocytes. Cryobiology. 2005 Aug;51(1):29–53.

27. Berrada MS, Bischof JC. Evaluation of freezing effects on human microvascular-endothelial cells (HMEC). Cryo Letters. 2001 Nov-Dec;22(6):353–66.

28. Stott SL, Karlsson JOM. Visualization of intracellular ice formation using high-speed video cryomicroscopy. Cryobiology. 2009 Feb;58(1):84–95.

29. Smith DJ, Schulte M, Bischof JC. The effect of dimethylsulfoxide on the water transport response of rat hepatocytes during freezing. J Biomech Eng. 1998 Oct;120(5):549–58.

30. Xu Y, Zhao G, Zhou X, Ding W, Shu Z, Gao D. Biotransport and intracellular ice formation phenomena in freezing human embryonic kidney cells (HEK293T). Cryobiology. 2014 Apr;68(2):294–302.

31. Lauterboeck L, Wolkers WF, Glasmacher B. Cryobiological parameters of multipotent stromal cells obtained from different sources. Cryobiology. 2017 Feb;74:93–102.

32. Huang Y, Dong Y, Gao B, Ma R, Gao FL, Shen L. Transmembrane water transport and intracellular ice of human umbilical vein endothelial cells during freezing. Biopreserv Biobank. 2022 Aug;20(4):311–316.

33. Pakhomov O, Shevchenko N, Chernobai N, Prokopiuk V, Yershov S, Bozhok G. Open-source hardware-and software-based cryomicroscopy system for investigation of phase transitions in cryobiological research. J Microsc. 2024 Feb;293(2):71–85.

34. Pazhayannur PV, Bischof JC. Measurement and simulation of water transport during freezing in mammalian liver tissue. J Biomech Eng. 1997 Aug;119(3):269–77.

35. Devireddy RV, Raha D, Bischof JC. Measurement of water transport during freezing in cell suspensions using a differential scanning calorimeter. Cryobiology. 1998 Mar;36(2):124–55.

36. Devireddy RV, Bischof JC. Measurement of water transport during freezing in mammalian liver tissue: Part II--The use of differential scanning calorimetry. J Biomech Eng. 1998 Oct;120(5):559–69.

37. Barratt PR, Devireddy RV, Storey KB, Bischof JC. Biophysics of freezing in liver of the freeze-tolerant wood frog, R. sylvatica. Ann N Y Acad Sci. 1998 Sep 11;858:284–97.

38. Devireddy RV, Barratt PR, Storey KB, Bischof JC. Liver freezing response of the freeze-tolerant wood frog, Rana sylvatica, in the presence and absence of glucose. I. Experimental measures. Cryobiology. 1999 Jun;38(4):310–26

39. Devireddy RV, Barratt PR, Storey KB, Bischof JC. Liver freezing response of the freeze-tolerant wood frog, Rana sylvatica, in the presence and absence of glucose. II. Mathematical modeling. Cryobiology. 1999 Jun;38(4):327–38.

40. Devireddy RV, Coad JE, Bischof JC. Microscopic and calorimetric assessment of freezing processes in uterine fibroid tumor tissue. Cryobiology. 2001 Jun;42(4):225–43.

41. Devireddy RV, Li G, Leibo SP. Suprazero cooling conditions significantly influence subzero permeability parameters of mammalian ovarian tissue. Mol Reprod Dev. 2006 Mar;73(3):330–41

42. Kardak A, Leibo SP, Devireddy R. Membrane transport properties of equine and macaque ovarian tissues frozen in mixtures of dimethylsulfoxide and ethylene glycol. J Biomech Eng. 2007 Oct;129(5):688–94.

43. Devireddy RV. Cryobiology of ovarian tissues: Known knowns and known unknowns. Minerva Ginecol. 2018 Aug;70(4):387–401.

44. Consiglio AN, Rubinsky B, Powell-Palm MJ. Thermodynamic analysis of a partial freezing organ preservation protocol. Cryobiology. 2026 Jun;123:105636.

45. McGrath JJ. Preservation of biological material by freezing and thawing. In: Shitzer A, Eberhart RC, editors. Heat transfer in medicine and biology: analysis and applications. Vol. 2. New York (NY): Plenum Press; 1985. p. 185–238.

46. Karlsson JO, Toner M. Long-term storage of tissues by cryopreservation: critical issues. Biomaterials. 1996;17(3):243–256.

47. Zhmakin AI. Fundamentals of cryobiology: 1. Introduction. 2. Ice formation in biological medium. 3. Biological effect of low temperatures. 4. Imitation models. 5. Microscopic models. 6. Macroscopic models. Conclusions. A. Brief history of cryomedicine. B. Simulation of solidification. C. Thermal properties of tissues. Berlin: Springer; 2008.

48. Whaley D, Damyar K, Witek R, Mendoza A, Alexander M, Lakey J. Cryopreservation: an overview of principles and cell-specific considerations. Cell Transplant. 2021;30:963689721999617.

49. Bischof JC. Quantitative measurement and prediction of biophysical response during freezing in tissues. Annu Rev Biomed Eng. 2000;2(1):257–288.

50. Steif PS, Palastro MC, Rabin Y. The effect of temperature gradients on stress development during cryopreservation via vitrification. Cell Preserv Technol. 2007;5(2):104–115.

51. Joshi P, Rabin Y. Thermomechanical stress analyses of nanowarming-assisted recovery from cryopreservation by vitrification in human heart and rat heart models. PLoS One. 2023;18(8):e0290063.

52. Karlsson JO. A theoretical model of intracellular devitrification. Cryobiology. 2001;42(3):154–169.

53. Kumano H, Asaoka T, Saito A, Okawa S. Study on latent heat of fusion of ice in aqueous solutions. International Journal of Refrigeration. 2007 Mar 1;30(2):267–73.

54. He X, Bischof JC. Analysis of thermal stress in cryosurgery of kidneys. J BiomechEng. 2005;127:656–661.

55. Rabin Y, Steif PS. Analysis of thermal stresses around a cryosurgical probe. Cryobiology. 1996;33(2):276–290.

56. Rabin Y, Steif PS, Taylor MJ, Julian TB, Wolmark N. An experimental study of the mechanical response of frozen biological tissues at cryogenic temperatures. Cryobiology. 1996;33(4):472–482.

57. Yoo J, Rubinsky B. Numerical computation using finite elements for the moving interface in heat transfer problems with phase transformation. Numerical Heat Transfer. 1983 Apr 1;6(2):209–22.

58. Li M, Chaouki H, Robert JL, Ziegler D, Fafard M. Numerical simulation of Stefan problem coupled with mass transport in a binary system through XFEM/level set method. Journal of Scientific Computing. 2019 Jan 15;78(1):145–66.

59. Körber C. Phenomena at the advancing ice–liquid interface: solutes, particles and biological cells. Quarterly reviews of biophysics. 1988 May;21(2):229–98.

60. Devireddy RV, Bischof, JC, Leo PH, Lowengrub JS. Measurement and numerical analysis of freezing in solutions enclosed in a small container. Int J Heat Mass Transf. 2002; 45(9):1915–1931.

61. Han B, Bischof JC. Thermodynamic nonequilibrium phase change behavior and thermal properties of biological solutions for cryobiology applications. J Biomech Eng. 2004 Apr;126(2):196–203.

62. Han B, Choi JH, Dantzig JA, Bischof JC. A quantitative analysis on latent heat of an aqueous binary mixture. Cryobiology. 2006 Feb;52(1):146–51.

63. Kumano H, Asaoka T, Saito A, Okawa S. Formulation of latent heat of fusion in aqueous solutions. Int J Ref. 2009; 32(1):175–182.

64. Johnson S, Hall C, Das S, Devireddy R. Freezing of Solute-Laden Aqueous Solutions: Kinetics of Crystallization and Heat- and Mass-Transfer-Limited Model. Bioengineering (Basel). 2022 Oct 10;9(10):540.

65. Rubinsky B. Solidification processes in saline solutions. J Cryst Growth. 1983;62:513–522.

66. Zhang A, Xu LX, Sandison GA, Zhang J. A microscale model for prediction of breast cancer cell damage during cryosurgery. Cryobiology. 2003 Oct;47(2):143–54.

67. Silva M, Freitas B, Andrade R, Espregueira-Mendes J, Silva F, Carvalho Ó, Flores P. Computational Modelling of the Bioheat Transfer Process in Human Skin Subjected to Direct Heating and/or Cooling Sources: A Systematic Review. Ann Biomed Eng. 2020 Jun;48(6):1616–1639.

68. Saeedi A, Devireddy R, Kothari M. A thermo-mechanically coupled finite-deformation model for freezing-induced damage in soft materials. J Mech Phys Solids. 2026;106545.

69. Rubinsky B, Pegg DE. A mathematical model for the freezing process in biological tissue. Proc R Soc Lond B Biol Sci. 1988;234:343-358.

70. Bischof JC, Rubinsky B. Microscale heat and mass transfer of vascular and intracellular freezing in the liver. ASME J Heat Transfer. 1993;115:1029–1035.

71. McGrath JJ. Quantitative measurement of cell membrane transport: technology and applications. Cryobiology. 1997 Jun;34(4):315–34.

72. Anderson DM, Benson JD, Kearsley AJ. Foundations of modeling in cryobiology-I: Concentration, Gibbs energy, and chemical potential relationships. Cryobiology. 2014 Dec;69(3):349–60.

73. Anderson DM, Benson JD, Kearsley AJ. Foundations of modeling in cryobiology-II: Heat and mass transport in bulk and at cell membrane and ice-liquid interfaces. Cryobiology. 2019 Dec;91:3–17.

74. Anderson DM, Benson JD, Kearsley AJ. Foundations of modeling in cryobiology-III: Inward solidification of a ternary solution towards a permeable spherical cell in the dilute limit. Cryobiology. 2020 Feb 1;92:34–46.

75. Benson JD. Mathematical Modeling and optimization of cryopreservation in single cells. Methods Mol Biol. 2021;2180:129–172.

76. Toner M, Cravalho EG, Karel M. Thermodynamics and kinetics of intracellular ice formation during freezing of biological cells. Journal of Applied Physics. 1990 Feb 1;67(3):1582–93.

77. Toner M, Tompkins RG, Cravalho EG, Yarmush ML. Transport phenomena during freezing of isolated hepatocytes. AIChE journal. 1992 Oct;38(10):1512–22.

78. Toner M. Nucleation of ice crystals in biological cells. In Advances in Low-Temperature Biology (Steponkus, P. L., Ed.), JAI Press, London 2, 1993; pp. 1–52.

79. Diller KR, Cravalho EG, Huggins CE. Intracellular freezing in biomaterials. Cryobiology. 1972 Oct;9(5):429–40.

80. Pitt RE, Steponkus PL. Quantitative analysis of the probability of intracellular ice formation during freezing of isolated protoplasts. Cryobiology. 1989 Feb;26(1):44–63.

81. Pitt RE, Chandrasekaran M, Parks JE. Performance of a kinetic model for intracellular ice formation based on the extent of supercooling. Cryobiology. 1992 Jun;29(3):359–73.

82. Acker JP, Larese A, Yang H, Petrenko A, McGann LE. Intracellular ice formation is affected by cell interactions. Cryobiology. 1999 Jun 1;38(4):363–71.

83. Irimia D, Karlsson JO. Kinetics of intracellular ice formation in one-dimensional arrays of interacting biological cells. Biophys J. 2005 Jan;88(1):647–60.

84. Li W, Yang G, Zhang A, Xu LX. Numerical study of cell cryo-preservation: a network model of intracellular ice formation. PLoS One. 2013;8(3):e58343.

85. Yi J, Liang XM, Zhao G, He X. An improved model for nucleation-limited ice formation in living cells during freezing. PLoS One. 2014 May 22;9(5):e98132. doi: 10.1371/journal.pone.0098132.

86. Amiri F, Benson JD. A three-dimensional lattice-free agent-based model of intracellular ice formation and propagation and intercellular mechanics in liver tissues. R Soc Open Sci. 2024 Jul 17;11(7):231337.

87. Franks F, Mathias SF, Galfre P, Webster SD, Brown D. Ice nucleation and freezing in undercooled cells. Cryobiology. 1983 Jun;20(3):298–309.

88. Devireddy RV, Smith DJ, Bischof JC. Mass transfer during freezing in rat prostate tumor tissue. AIChE J. 1999;45(3):639–654.

89. Patankar S. Numerical heat transfer and fluid flow. Taylor & Francis; 2018 Oct 8.

90. Junkins JL, Bani Younes A, Woollands RM, Bai X. Picard iteration, chebyshev polynomials and chebyshev-picard methods: Application in astrodynamics. The Journal of the Astronautical Sciences. 2013 Dec;60(3):623–53.

91. Woollands RM, Bani Younes A, Junkins JL. New solutions for the perturbed lambert problem using regularization and picard iteration. Journal of Guidance, Control, and Dynamics. 2015 Sep;38(9):1548–62.

92. P Kunze HE, Vrscay ER. Solving inverse problems for ordinary differential equations using the Picard contraction mapping. Inverse Problems. 1999 Jun 1;15(3):745–70.

93. Gao C, Wang Y. A general formulation of Peaceman and Rachford ADI method for the N-dimensional heat diffusion equation. International communications in heat and mass transfer. 1996 Oct 1;23(6):845–54.

94. Dai W, Nassar R. Compact ADI method for solving parabolic differential equations. Numerical Methods for Partial Differential Equations: An International Journal. 2002 Mar;18(2):129–42.

95. Schrage DS. On the application of ADI methods to predict conjugate phase change and diffusion heat transfer. Numerical Heat Transfer: Part B: Fundamentals. 2001 Jun 1;39(6):563–83.

96. Albu AF, Zubov VI. Identification of the thermal conductivity coefficient in the three-dimensional case by solving a corresponding optimization problem. Computational Mathematics and Mathematical Physics. 2021 Sep;61(9):1416–31.

97. Yang J, Xie Z, Meng HJ, Liu WH, Ji ZP. Multiple time steps optimization for real-time heat transfer model of continuous casting billets. International Journal of Heat and Mass Transfer. 2014 Sep 1;76:492–8.

98. Qing-Cheng W, Zhao-Chun W, Xiang-Ping Z. The study of the temperature field in an infinite slab under line and plane heat source. International Journal of Numerical Methods for Heat & Fluid Flow. 2015 Jan 5;25(1):25–32.

99. Zeng P, Deng ZS, Liu J. Parallel algorithms for freezing problems during cryosurgery. International Journal of Information Engineering and Electronic Business. 2011 Mar 1;3(2):11.

100. Zhu He Z, Xue X, Liu J. An effective finite difference method for simulation of bioheat transfer in irregular tissues. Journal of heat transfer. 2013 Jul 1;135(7):071003.

101. Morley MJ, Fursey GA. The apparent specific heat and enthalpy of fatty tissue during cooling. International Journal of Food Science & Technology. 1988 Oct;23(5):467–77.

102. Kundu P, Sukumar S, Kar SP. Numerical modeling for freezing and cryogenic preservation for viability of biological tissue. Materials Today: Proceedings. 2018 Jan 1;5(9):18823–32.

103. Pavlyuk EV. Mathematical modeling of conductive–convective melting with a two-phase region by the enthalpy-porosity method. The European Physical Journal Special Topics. 2024 Dec;233(23):3321–33.

104. Andrushkiw R. Mathematical modeling of freezing front propagation in biological tissue. Mathematical and Computer Modelling. 1990 Jan 1;13(10):1–9.

105. Olien CR, Livingston III DP. Understanding freeze stress in biological tissues: Thermodynamics of interfacial water. Thermochimica Acta. 2006 Dec 1;451(1-2):52–6.

106. Majchrzak E, Ladyga E. Numerical analysis of freezing of a binary solution during cryosurgical process using the Boundary Element Method. WIT Transactions on Modelling and Simulation. 1997 Jul 30;18.

107. Dombrovsky LA, Nenarokomova NB, Tsiganov DI, Zeigarnik YA. Modeling of repeating freezing of biological tissues and analysis of possible microwave monitoring of local regions of thawing. International Journal of Heat and Mass Transfer. 2015 Oct 1;89:894–902.

108. Lee CY, Bastacky J. Comparative mathematical analyses of freezing in lung and solid tissue. Cryobiology. 1995 Aug 1;32(4):299–305.

109. Kumar S, Katiyar VK. Mathematical modeling of freezing and thawing process in tissues: A porous media approach. International Journal of Applied Mechanics. 2010 Sep;2(03):617–33.

110. Bakhach J. The cryopreservation of composite tissues: Principles and recent advancement on cryopreservation of different type of tissues. Organogenesis. 2009 Jul 1;5(3):119–26.

111. Li FF, Liu J, Yue K. Exact analytical solution to three-dimensional phase change heat transfer problems in biological tissues subject to freezing. Applied Mathematics and Mechanics. 2009 Jan;30(1):63–72.

112. Wills VA, McCartney DG. Modelling of dendritic solidification using finite element method. Materials Science and Technology. 1992 Feb 1;8(2):114–22.

113. Kurtz W, Fisher D J. Fundamentals of Solidification. Switzerland: Trans Tech SA, 1984;242.

114. Cantor B. Fundamentals of rapid solidification. Science and Technology of the Undercooled Melt: Rapid Solidification Materials and Technologies. 1986:3–28.

115. Wu M, Ludwig A, Kharicha A. Volume-averaged modeling of multiphase flow phenomena during alloy solidification. Metals. 2019 Feb 14;9(2):229.

116. Bouchard D, Kirkaldy JS. Scaling of intra-granular dendritic microstructure in ingot solidification. Metallurgical and Materials Transactions B. 1996 Feb;27(1):101–13.

117. Kumar S, Katiyar VK. Transient analysis of alloy freezing in finite media with energy generation and convective cooling. International Journal of Applied Mechanics and Engineering. 2010;15(4):1155–68.

118. Babu SS. Thermodynamic and kinetic models for describing microstructure evolution during joining of metals and alloys. International Materials Reviews. 2009 Nov 1;54(6):333–67.

119. Chang T, Zhao G. Ice inhibition for cryopreservation: materials, strategies, and challenges. Advanced Science. 2021 Mar;8(6):2002425.

120. Li B, Sun DW. Novel methods for rapid freezing and thawing of foods–a review. Journal of food engineering. 2002 Sep 1;54(3):175–82.

121. Huo Y, Rao Z. The quasi-enthalpy based lattice Boltzmann model for solid-liquid phase change. Applied Thermal Engineering. 2017 Mar 25;115:1237–44.

122. Cogné C, Nguyen PU, Lanoisellé JL, Van Hecke E, Clausse D. Modeling heat and mass transfer during vacuum freezing of puree droplet. international journal of refrigeration. 2013 Jun 1;36(4):1319–26.

123. Hoang DK, Lovatt SJ, Olatunji JR, Carson JK. Validated numerical model of heat transfer in the forced air freezing of bulk packed whole chickens. International Journal of Refrigeration. 2020 Oct 1;118:93–103.

124. Alabi KP, Zhu Z, Sun DW. Transport phenomena and their effect on microstructure of frozen fruits and vegetables. Trends in Food Science & Technology. 2020 Jul 1;101:63–72.

125. Pham QT. Modelling heat and mass transfer in frozen foods: A review. International Journal of Refrigeration. 2006 Sep 1;29(6):876–88.

126. Zhang G, You J, Yang H. Effect of freezing methods on the physical and chemical properties and protein oxidation of porcine liver. Science and Technology of Food Industry. 2025 Apr 10;46(3):342–9.

127. Gangwar L, Han Z, Scheithauer C, Namsrai BE, Kantesaria S, Goldstein R, Etheridge ML, Finger EB, Bischof JC. Physical vitrification and nanowarming at liter-scale CPA volumes: toward organ cryopreservation. Nat Commun. 2025 Sep 26;16(1):8511.

128. McGann LE. Differing actions of penetrating and nonpenetrating cryoprotective agents. Cryobiology. 1978 Aug;15(4):382–90.

129. Ahmadkhani N, Sugden C, Benson JD, Eroglu A, Higgins AZ. High-throughput evaluation of cryoprotective agents for mixture effects that reduce toxicity. Cryobiology. 2025 Dec;121:105316.

130. Kedem O, Katchalsky A. Thermodynamic analysis of the permeability of biological membranes to non-electrolytes. Biochim Biophys Acta. 1958 Feb;27(2):229–46.

131. Fuhrman FA. Transport through biological membranes. Annu Rev Physiol. 1959;21:19–48.

132. Kedem O, Essig A. Isotope flows and flux ratios in biological membranes. J Gen Physiol. 1965 Jul;48(6):1047–70.

133. Slezak A, Turczynski B. Modification of the Kedem-Katchalsky equations. Biophys Chem. 1986 Jul;24(2):173–8.

134. Hoshiko T, Lindley BD. Phenomenological description of active transport of salt and water. J Gen Physiol. 1967 Jan;50(3):729–58.

135. Kleinhans FW. Membrane permeability modeling: Kedem-Katchalsky vs a two-parameter formalism. Cryobiology. 1998 Dec;37(4):271–89.

136. Suchanek G. On the derivation of the Kargol’s mechanistic transport equations from the Kedem-Katchalsky phenomenological equations. Gen Physiol Biophys. 2005 Jun;24(2):247–58.

137. Devireddy RV, Amorim CA, Leibo SP. Permeability characteristics of ovine primordial follicles calculated with two parameter Kedem-Katchalsky formulation. Cell Preservation Technology. 2006 Sep 1;4(3):188–98.

138. He Y, Devireddy RV. An inverse approach to determine solute and solvent permeability parameters in artificial tissues. Ann Biomed Eng. 2005 May;33(5):709–18.

139. Levin RL. A generalized method for the minimization of cellular osmotic stresses and strains during the introduction and removal of permeable cryoprotectants. J Biomech Eng. 1982 May;104(2):81–6.

140. Nei T. Mechanism of hemolysis by freezing at near zero temperatures. II. Investigation of factors affecting hemolysis by freezing. Cryobiology. 1967;4:303–308.

141. Ishiguro H, Rubinsky B. Mechanical interactions between ice crystals and red bloodcells during directional solidification. Cryobiology. 1994;31:483–500.

142. Li X, Zhang S, Zhang Y, Zhou X. Visualization of intracellular ice formation and growth in mouse oocytes. Cryo Letters. 2024 May-Jun;45(3):185–193.

143. Karlsson JO. Theoretical analysis of unidirectional intercellular ice propagation in stratified cell clusters. Cryobiology. 2004 Jun;48(3):357–61

144. Xu F, Moon S, Zhang X, Shao L, Song YS, Demirci U. Multi-scale heat and mass transfer modelling of cell and tissue cryopreservation. Philos Trans A Math Phys Eng Sci. 2010 Feb 13;368(1912):561–83.

145. Kumar S. Numerical analysis of phase-change heat transfer in axisymmetric biological tissue during nano-cryosurgery. Heat Transfer. 2023 Sep;52(6):4274–92.

146. Bellil M, Saidane A, Bennaoum M. A TLM study of bioheat transfer during freeze-thaw cryosurgery. Biomedical Physics & Engineering Express. 2018 Nov 1;4(6):065031.

147. Bao Y, Liu H, Tang J, Du W. A systematic review of bioheat transfer models in cryosurgery. Cryogenics. 2026 Jan 29:104295.

148. Tang J, Bao Y, Du W, Wei W. Numerical simulation of bioheat transfer during bronchial cryobiopsy using cryoprobes of different diameters. Cryobiology. 2025 Dec 1;121:105320.

149. Liu Y, He L, Zhao G. Multiscale heat and mass transfer in biomaterial cryopreservation. International Journal of Heat and Mass Transfer. 2025 Dec 1;252:127414.

150. Bai B, Xue C, Wen Y, Lim J, Le Z, Shou Y, Shin S, Tay A. Cryopreservation in the era of cell therapy: revisiting fundamental concepts to enable future technologies. Advanced Functional Materials. 2023 Oct;33(40):230337

151. Tang X, Pikal MJ. Design of freeze-drying processes for pharmaceuticals: practical advice. Pharmaceutical research. 2004 Feb;21(2):191–200.

152. Hottot A, Peczalski R, Vessot S, Andrieu J. Freeze-drying of pharmaceutical proteins in vials: modeling of freezing and sublimation steps. Drying Technology. 2006 Jun 1;24(5):561–70.

153. Bhatnagar B, Tchessalov S. Advances in freeze drying of biologics and future challenges and opportunities. Drying Technologies for Biotechnology and Pharmaceutical Applications. 2020 Apr 6:137–77.

154. Adams GD, Cook I, Ward KR. The principles of freeze-drying. In Cryopreservation and freeze-drying protocols 2014 Nov 14 (pp. 121–143). New York, NY: Springer New York.

155. Abla KK, Mehanna MM. Freeze-drying: A flourishing strategy to fabricate stable pharmaceutical and biological products. International Journal of Pharmaceutics. 2022 Nov 25;628:122233.

156. Jakubowska E, Bielejewski M, Milanowski B, Lulek J. Freeze-drying of drug nanosuspension–study of formulation and processing factors for the optimization and characterization of redispersible cilostazol nanocrystals. Journal of Drug Delivery Science and Technology. 2022 Aug 1;74:103528.

